# Secretin channel-interactors prevent antibiotic influx during type IV pili assembly in *Pseudomonas aeruginosa*

**DOI:** 10.1101/2021.09.13.460190

**Authors:** Oh Hyun Kwon, Joel W Sher, Bi-o Kim, You-Hee Cho, Hongbaek Cho

**Affiliations:** Department of Biological Sciences, College of Natural Sciences, Sungkyunkwan University, Republic of Korea; Department of Microbiology, Harvard Medical School, Boston, United States; Department of Pharmacy, College of Pharmacy and Institute of Pharmaceutical Sciences CHA University, Republic of Korea

**Author notes:** To whom correspondence should be addressed. Hongbaek Cho, Ph.D., Sungkyunkwan University Department of Biological Sciences Suwon 16419, Republic of Korea.

**Keywords:** type IV pili, secretin, outer membrane, permeability barrier, antibiotics, drug resistance, macrolide

## Abstract

Type IV pili (T4P) are important virulence factors involved in host attachment and other aspects of bacterial pathogenesis. In Gram-negative bacteria, the T4P filament is polymerized from pilin subunits at the platform complex in the inner membrane (IM) and exits the outer membrane (OM) through the OM secretin channel. Although essential for T4P assembly and function, the OM secretin complexes can potentially impair the permeability barrier function of the OM and allow the entry of antibiotics and other toxic molecules. The mechanism by which Gram-negative bacteria prevent secretin-mediated OM leakage is currently not well understood. Here, we report a discovery of SlkA and SlkB (PA5122 and PA5123) that prevent permeation of several classes of antibiotics through the secretin channel of *Pseudomonas aeruginosa* type IV pili. We found these periplasmic proteins interact with the OM secretin complex and prevent toxic molecules from entering through the channel when there is a problem in the assembly of the T4P IM subcomplexes or when docking between the OM and IM complexes is defective. Thus, our results indicate that the secretin channel-interacting proteins play an important role in maintaining the OM permeability barrier, suggesting they may be attractive targets for potentiators that sensitize Gram-negative pathogens to antibiotics that are normally ineffective at penetrating the OM.

## Introduction

Gram-negative bacterial pathogens such as *Pseudomonas aeruginosa*, *Acinetobacter baumannii*, and *Enterobacteriaceae* are often multidrug-resistant because their cell envelope functions as an effective permeability barrier against a variety of antibiotics (Nikaido, 2003; Delcour, 2009). Their cell envelope is a multi-layered structure consisting of the cytoplasmic (inner) membrane (IM), the outer membrane (OM), and the peptidoglycan layer in the periplasmic space between the two membranes. Among these layers, the OM plays a crucial role in the barrier function of the envelope due to its asymmetric bilayer structure consisting of two types of membrane lipids: lipopolysaccharides (LPS) in the outer leaflet and phospholipids in the inner leaflet. The long glycan chains of the LPS limits the diffusion of hydrophobic compounds and the lipid bilayer prevents the entry of hydrophilic compounds across the OM. In addition, RND type efflux pumps that span the entire envelope efficiently remove various toxic molecules from both the cytoplasm and periplasm, reinforcing the barrier function of the OM (Silver, 2011; Raetz & Whitfield, 2002; Krishnamoorthy *et al*, 2017; Li *et al*, 2015).

The permeability barrier formed by the OM is necessarily imperfect because it must allow the entry of nutrients required for bacterial growth. To promote diffusion across the OM, Gram-negative bacteria assemble a variety of integral outer membrane proteins (OMPs) to transport molecules across the OM and the pore structure of these OMPs compromise the permeability barrier. General porins and substrate-specific channels responsible for nutrient transport have been shown to facilitate the diffusion of small hydrophilic compounds into the periplasm through their open pore structure with hydrophilic inner rim (Pagès *et al*, 2008; Prajapati *et al*, 2021). In *Enterobacteriaceae*, loss of the general porin OmpF results in elevated resistance to small hydrophilic antibiotics such as beta-lactams (Delcour, 2009; Sugawara *et al*, 2016; Choi & Lee, 2019; Vergalli *et al*, 2020). In *P. aeruginosa*, OprD, which is normally responsible for the transport of basic amino acids and peptides, functions as a carbapenem entry channel and an *oprD* mutation is frequently observed in carbapenem-resistant clinical isolates (Fukuoka *et al*, 1993; Lee & Ko, 2012; Kim *et al*, 2016; Wolter *et al*, 2004).

Besides the OMPs whose pore structure is formed from a single polypeptide, Gram-negative pathogens also assemble multimeric channels in the OM as part of large transenvelope complexes like pili and secretion systems, many of which are critical for virulence and survival in the host (Nikaido, 2003). A group of homologous proteins called secretins form 12- to 15-meric OM channel complexes to accommodate protein substrates in several envelope-spanning virulence systems: type II secretion systems (T2SS), type III secretion systems (T3SS), and type IV pili assembly systems (T4PS) (Majewski *et al*, 2018; Korotkov *et al*, 2011). Cryo-electron microscopy (cryo-EM) of the secretin complexes revealed that these proteins share a similar architecture. A less-conserved N-terminal domain forms a periplasmic vestibule that interacts with the IM components of the virulence system, and a well-conserved C-terminal domain forms an OM channel with one or two gate structures (Weaver *et al*, 2020; Hu *et al*, 2018; D’Imprima *et al*, 2017; Naskar *et al*, 2021). Although secretin channels form much larger pore structures than the channels formed by a single polypeptide, it has been assumed that the gate structures of secretins prevent leakage of periplasmic contents and permeation of extracellular chemicals when they are not in use for protein secretion or pilus assembly (Majewski *et al*, 2018; Korotkov *et al*, 2011). However, *in vitro* studies have suggested that secretin channels are not completely sealed in their resting state and that diffusion of small compounds occurs through these channels (Disconzi *et al*, 2014). In addition, mutations in the secretin gene *pilQ* of T4P have been reported to increase the MIC of several antibiotics in pathogenic *Neisseria* species (Tzeng *et al*, 2019; Zhao *et al*, 2005; Nandi *et al*, 2015). These results indicate that the secretin complex can function as a conduit that allows the entry of antibiotics across the OM although mechanistic details of drug entry have remained largely unclear.

To uncover factors important for maintaining the OM permeability barrier in *P. aeruginosa*, we employed transposon sequencing (Tn-seq). A transposon mutant library was challenged with the hydrophobic antibiotic erythromycin at a concentration that normally does not cause a growth defect due to the barrier function of the OM. Comparison of the transposon profiles between treated and untreated samples identified two orthologous genes *PA5122* and *PA5123*, encoding putative periplasmic proteins that are required for normal erythromycin resistance. We discovered that these proteins prevent <u>s</u>ecretin lea<u>k</u>iness and therefore have renamed them SlkA and SlkB (pronounced “slick-A” and “slick-B”). Genetic and microscopic analyses suggested that the Slk proteins interact with the secretin complex of the T4PS to prevent the diffusion of drugs across the OM when the OM secretin complex is not docked with the IM complex. Overall, our results indicate that the gate structure found in secretin channels is insufficient to maintain the OM permeability barrier and that additional partner proteins are required to prevent the entry of toxic compounds during the vulnerable stages of T4P assembly.

## Results

### PA5122 and PA5123 (SlkA and SlkB) are periplasmic proteins involved in maintenance of permeability barrier function

To identify factors important for maintaining OM barrier function in *P. aeruginosa* against hydrophobic antibiotics, we employed a Tn-seq approach with erythromycin treatment (Fig. 1A). Wild-type *P. aeruginosa* PAO1 cells were mutagenized by random insertion of a mariner transposon from pBTK30 that was transferred via conjugation. The resulting mutant library was grown for roughly ten generations in LB lacking or containing 10 μg/mL of erythromycin, which normally does not cause a growth defect in *P. aeruginosa* due to the permeability barrier function of the OM and the activity of RND efflux pumps. Comparison of the transposon insertion profile between erythromycin-treated and untreated samples revealed marked reduction of transposon insertions in *oprM* and *PA5122* in the drug-treated sample, which suggested that these genes play important roles for survival under erythromycin treatment (Fig. 1B). OprM is an outer membrane channel component of major RND efflux systems MexAB-OprM and MexXY-OprM that are responsible for extrusion of various toxic compounds and antibiotics from both periplasm and cytoplasm (Li *et al*, 2015, 1995; Tsutsumi *et al*, 2019). Thus, the *oprM* mutant is likely to be hypersensitive to erythromycin due to defective efflux activity. The function of PA5122 has not been documented yet. We therefore focused our efforts on revealing its cellular function.

**Figure 1.**
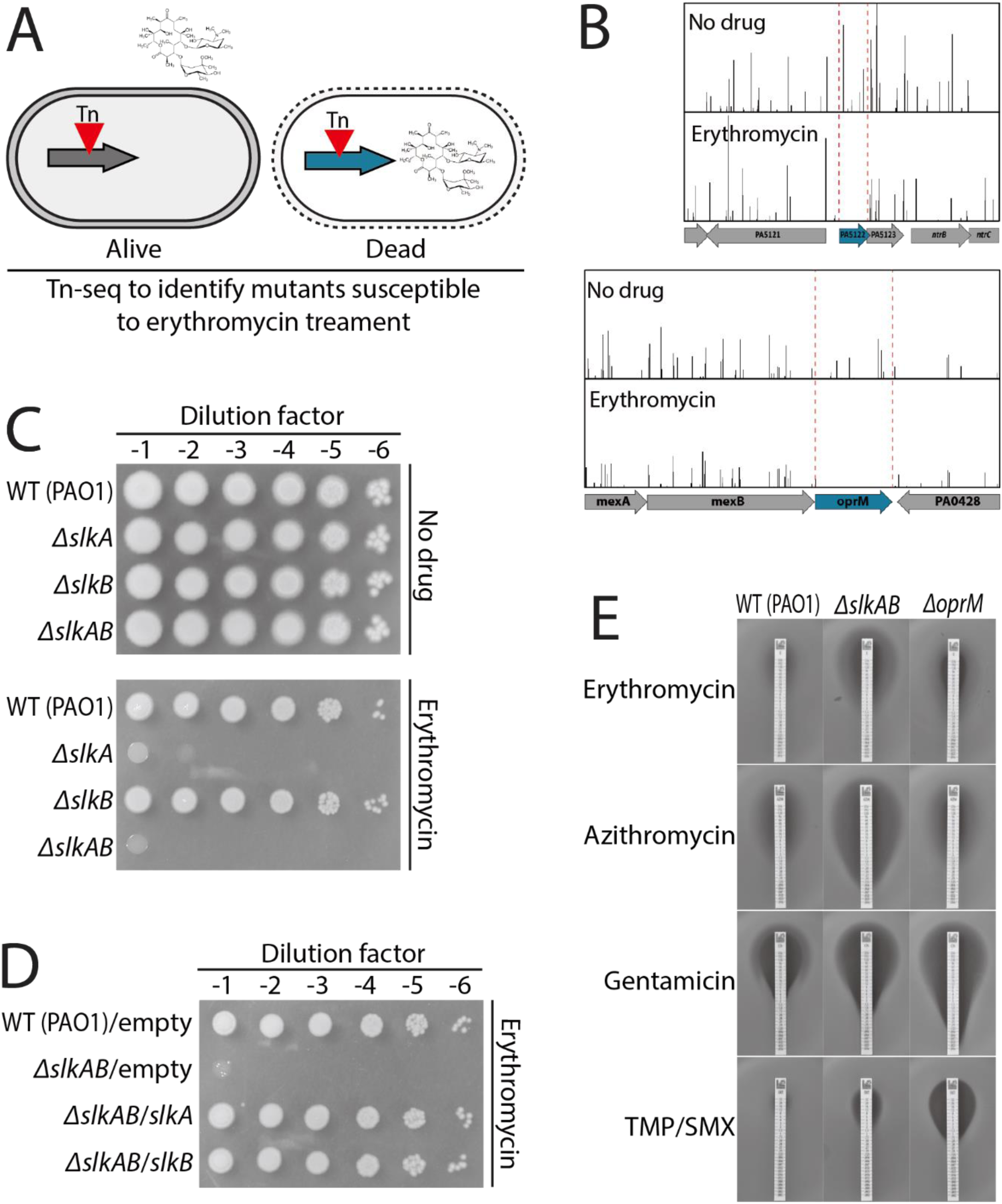
SlkA and SlkB play an important role in maintaining the envelope permeability barrier. (A) Scheme for identifying mutations that cause an envelope barrier defect. Mutants that have a defect in the permeability barrier function become susceptible to erythromycin treatment and mutations that cause erythromycin susceptibility are identified by a Tn-seq approach. (B) Transposon insertion profiles for *oprM* and *PA5122-PA5123* (*slkAB*) loci. Lines above the locus map represent transposon insertion sites and the height of the lines reflects the number of sequencing reads at each site. The regions that show significantly fewer reads in the erythromycin-treated sample compared with the untreated sample are indicated with red dotted lines. (C) Spot dilution assay for PAO1 (WT), OKP5 (*ΔslkA*), OKP6 (*ΔslkB*), and OKP7 (*ΔslkAB*) strains on LB agar only or supplemented with 50 μg/mL erythromycin. The strains were grown overnight in LB at 37 °C and normalized for cell density (OD_600_ = 2). Normalized cultures were then serially diluted 10-fold and 5 μL of each dilution (10^-1^-10^-6^) was spotted onto LB agar lacking or containing 50 μg/mL erythromycin. The plates were incubated at 37 °C and photographed after 20 hours. (D) Spot dilution assay for PAO1 harboring pJN105 (empty vector) and OKP7 strains harboring pJN105, pOKP17 (*slkA*), or pOKP18 (*slkB*) on LB agar supplemented with 50 μg/mL erythromycin. (E) Comparison of the sensitivity of *ΔslkAB* and *ΔoprM* strains to various antibiotics. Cells of the PAO1, OKP7 (*ΔslkAB*), and OKP62 (*ΔoprM*) strains were mixed with molten soft agar and overlaid on LB plates. Test strips impregnated with indicated antibiotics in a concentration gradient were then placed on the lawn of cells and the plates were incubated for 24 hours at 37 °C before being photographed.

PA5122 is predicted to encode a periplasmic protein, suggesting that the function of PA5122 is likely to be related to maintaining the envelope barrier rather than inhibiting erythromycin action on the ribosome. PA5122 constitutes an operon with a downstream gene PA5123, which also encodes a putative periplasmic protein. Interestingly, PA5122 and PA5123 encode homologous proteins with 35.5% sequence identity and 50.8% similarity, suggesting that the cellular role of PA5123 might be similar to that of PA5122. Thus, we made individual mutants and a mutant deleted for both genes, and compared their susceptibility to erythromycin with a wild-type strain. In accordance with the Tn-seq data, the *ΔPA5122* and the *ΔPA5122-5123* strains showed erythromycin susceptibility, but the *ΔPA5123* strain did not (Fig. 1C). To see if this difference arises due to a polar effect of *PA5122* mutation on *PA5123* expression, we individually expressed *PA5122* and *PA5123* in the *ΔPA5122-5123* strain and tested for the suppression of erythromycin susceptibility. Expression of either *PA5122* or *PA5123* suppressed the erythromycin susceptibility (Fig. 1D), suggesting that PA5123 has a similar function to PA5122 in maintaining the permeability barrier of the *P. aeruginosa* envelope. Based on our study presented below, we discovered that PA5122 and PA5123 prevent chemical influx through the T4P secretin channel and will henceforth refer to these proteins as SlkA and SlkB for prevention of <u>s</u>ecretin lea<u>k</u>iness.

To test if inactivation of SlkA and SlkB causes a general defect in envelope barrier function, we examined the sensitivity of this mutant to various classes of antibiotics. The *ΔslkAB* strain showed increased sensitivity to several antibiotics of the macrolide and aminoglycoside classes, such as azithromycin, gentamicin, and tobramycin when compared with the parental PAO1 strain (Fig. 1E and Fig. S1). It also showed increased sensitivity to trimethoprim (TMP)-sulfamethoxazole (SMX). As macrolides are hydrophobic and large, we also tested if the *ΔslkAB* strain becomes generally sensitive to other lipophilic or bulky antibiotics such as rifampicin, fusidic acid, and vancomycin, but inactivation of SlkA and SlkB did not noticeably increases the sensitivity to these drugs (Fig. S1).

We also compared the antibiotic sensitivity of the *ΔslkAB* strain with that of the *ΔoprM* strain to test if the function of Slk proteins is related to the MexAB-OprM and MexXY-OprM efflux pumps. However, the antibiotic sensitivity profile of the *ΔslkAB* strain was quite different from that of the *ΔoprM* strain. The *ΔslkAB* strain was much more susceptible to macrolides than the *ΔoprM* strain, whereas the *ΔoprM* strain showed higher sensitivity to aminoglycosides and TMP/SMT than the *ΔslkAB* strain. In addition, the MIC of beta-lactam drugs did not change much in the *ΔslkAB* strain but was increased in the *ΔoprM* strain (Fig. S1). The difference in the antibiotic sensitivity profile between the *ΔslkAB* and *ΔoprM* strains suggested that Slk proteins are likely to have a cellular function independent of the MexAB-OprM and MexXY-OprM efflux pumps.

### Slk proteins prevent permeation of antibiotics through the OM secretin complex of the type IV pili

To gain insight into the cellular function of the Slk proteins, we performed a genetic selection for suppressors of the erythromycin sensitivity phenotype of mutants lacking these proteins. The *ΔslkAB* strain was mutagenized by random insertion of a mariner transposon, and the resulting mutant library was selected on LB plates containing 50 μg/mL erythromycin to obtain suppressors. Then, transposon insertion sites of thirteen suppressors were mapped by arbitrarily primed PCR. Among them, nine had insertions in *pilQ*, two had insertions upstream of *pilQ*, and one each were mapped in *pilM* and *pilP* (Fig. 2A). For the remaining suppressors, one had an insertion in *pilF* and the other was mapped 124 bases upstream of the *fimV* coding sequence. PilQ is a component of *P. aeruginosa* T4P that assembles into a secretin channel in the OM through which a pilus exits the cell. Mutations upstream of *pilQ* were reported to significantly lower PilQ protein levels and inhibit PilQ multimer assembly (Ayers *et al*, 2009). PilF is an OM lipoprotein known as a pilotin that promotes localization and multimer assembly of the secretin PilQ in the OM (Koo *et al*, 2008). FimV is an IM protein that is also required for PilQ secretin assembly in the OM (Wehbi *et al*, 2011). Overall, the identified suppressors all had mutations that inhibit assembly of the secretin complex of the T4P. Thus, assembly of the T4P secretin complex in the absence of Slk proteins seemed responsible for the susceptibility of *ΔslkAB* mutants to erythromycin. Indeed, erythromycin sensitivity of the *ΔslkAB* strain was suppressed when we introduced a *pilQ* null mutation, and the suppression was abrogated when *pilQ* was expressed from an ectopic site on the chromosome (the Tn7 integration locus) in the *ΔpilQ ΔslkAB* strain (Fig. 2B). Deletion of *pilMNOPQ* also suppressed the erythromycin sensitivity of the *ΔslkAB* strain, but *pilQ* expression alone was sufficient for abolishing the suppression, indicating that mutations upstream of *pilQ* suppressed the erythromycin sensitivity of the *ΔslkAB* strain via a polar effect on *pilQ* expression. Although a *pilQ* mutation suppressed macrolide sensitivity of the *ΔslkAB* strain, it did not increase the resistance of the wild-type PAO1 strain to macrolides (Fig. 2C and Fig. S2), suggesting that assembly of the secretin channel does not cause a permeability defect with normal expression of the *slkAB* genes.

**Figure 2.**
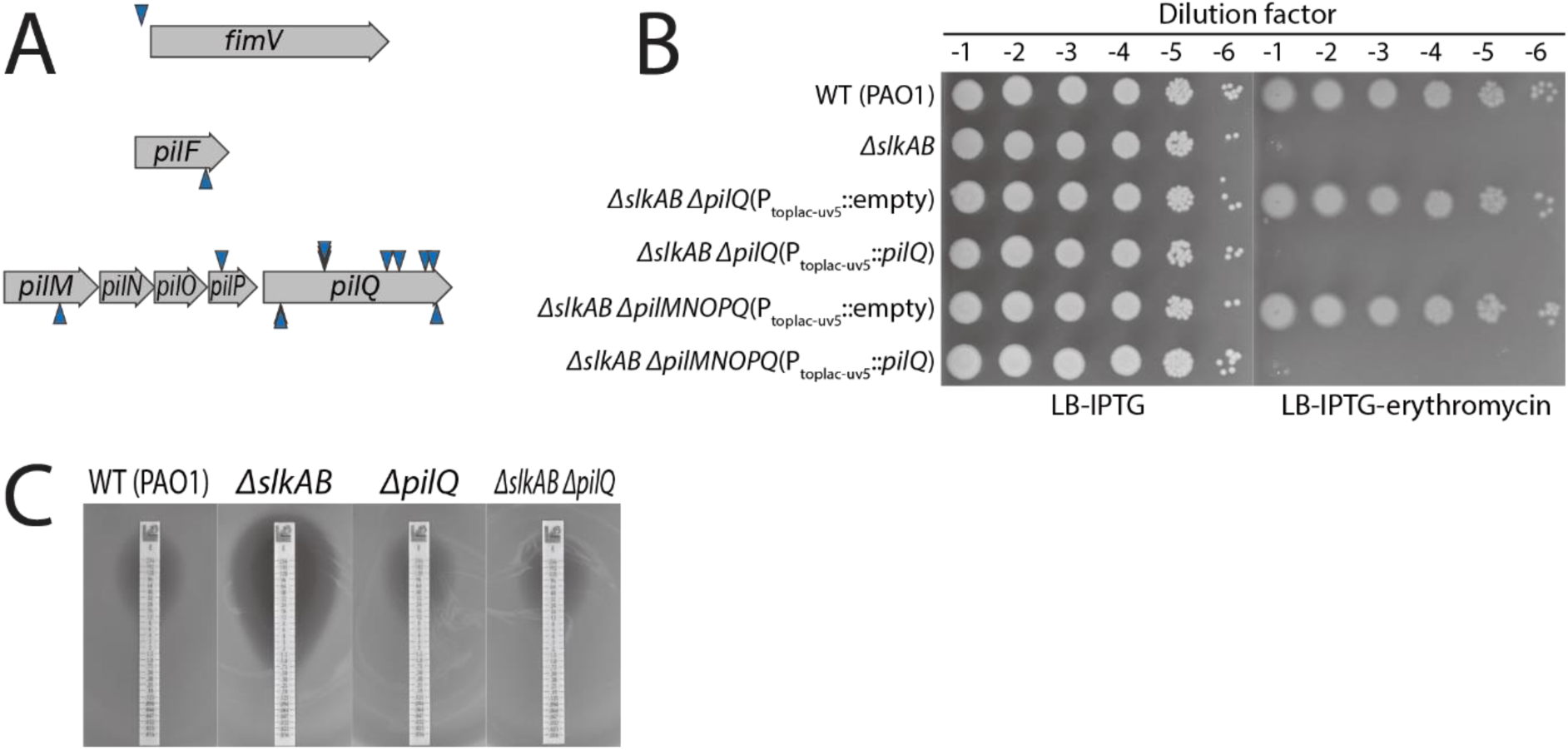
Assembly of the PilQ secretin complex of the T4PS causes the envelope barrier defect of the *ΔslkAB* strain. (A) A diagram of the transposon insertion sites of the suppressors isolated from the selection of the OKP7(*ΔslkAB*) strain on LB agar containing 50 μg/mL erythromycin. The triangles above the locus map represent the insertion for which the direction of transcription in the gentamicin resistance cassette of the transposon is in the same orientation as the disrupted gene; the triangles below, opposite direction. (B) OKP15 (*ΔslkAB ΔpilQ*) and OKP50 (*ΔslkAB ΔpilMNOPQ*) strains harboring a chromosomally integrated expression construct attTn7::pOKP121 [P_toplac-uv5_::*pilQ*] or an empty vector control attTn7::pKHT105 were grown in LB containing 1mM IPTG, serially diluted, and spotted onto indicated LB agar supplemented with 1mM IPTG to keep inducing *pilQ* expression. PAO1 and OKP7 (*ΔslkAB*) strains were also grown and spotted for comparison of growth phenotype. (C) Erythromycin sensitivity of PAO1, OKP7 (*ΔslkAB*), OKP14 (*ΔpilQ*), OKP15 (*ΔslkAB ΔpilQ*) strains was compared using erythromycin test strips as in Fig. 1E.

### Incomplete assembly of the T4P causes antibiotic permeation through the secretin channel in the absence of SlkA and SlkB

As *pilQ* deletion suppressed the erythromycin susceptibility of the *ΔslkAB* strain, we wondered if Slk proteins were previously unidentified components of the T4P and tested if they were required for T4P-driven twitching motility. Unexpectedly, even the wild type PAO1 strain used for the Tn-seq experiment and the initial characterization of the SlkAB phenotype was twitching-defective (Fig. 3A). As genetic variability among PAO1 strains has been widely recognized (Klockgether *et al*, 2009; Sidorenko *et al*, 2017; Chandler *et al*, 2019), we obtained a twitching-proficient PAO1 strain to examine phenotypes related to twitching motility. The twitching-proficient PAO1 strain will be referred to as PAO1^tw^ henceforth to distinguish it from the twitching-defective PAO1 strain. We introduced a *slkAB* mutation in several twitching-proficient *P. aeruginosa* strains PAO1^tw^, PA14, and PAK, and compared the twitching motility and erythromycin sensitivity between the WT strains and their *ΔslkAB* derivatives. A *slkAB* deletion did not cause an obvious defect of the twitching motility in any of the *P. aeruginosa* strains, showing that Slk proteins are not essential components of T4P (Fig. 3A). Surprisingly, however, deletion of *slkAB* did not cause erythromycin sensitivity in the twitching-proficient strains unlike in the twitching-defective PAO1 strain (Fig. 3B and Fig. S3). Although puzzling at first, this result led us to hypothesize that the erythromycin sensitivity of *slkAB* mutants might be related to the twitching motility defect of our original parent strain.

**Figure 3.**
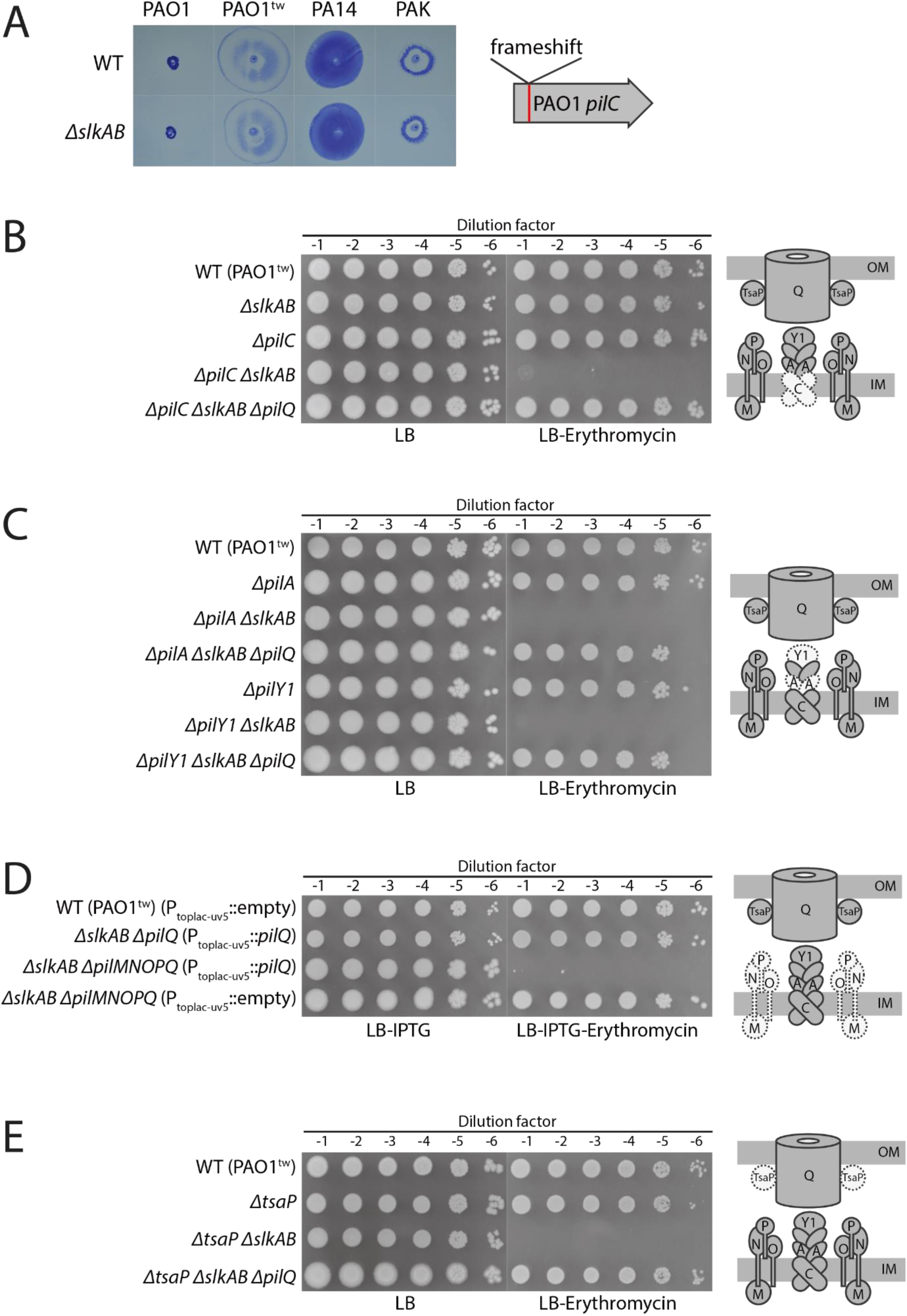
Effect of type IV pili mutations on the drug diffusion through the PilQ complex in the *ΔslkAB* strain. (A) Comparison of twitching motility among PAO1, PAO1^tw^, PA14, PAK, and their *ΔslkAB* derivatives. A single colony of each strain was stab-inoculated through LB agar to the polystyrene dish. After incubation for 48 hours at 30 °C, LB agar was removed and the cells attached to the polystyrene dish were stained with crystal violet and photographed. The diagram on the right shows a frameshift mutation in *pilC* of the PAO1 strain used for the Tn-seq and initial characterization. (B-E) Spot dilution assays to test the effect of T4P gene mutations on the drug permeation through the PilQ secretin complex of the *ΔslkAB* strain. Mutations in the platform (*pilC*), the pilus shaft (*pilA* and *pilY1*), alignment complex (*pilMNOP*), and *tsaP* genes were introduced in the PAO1^tw^ strain and its *ΔslkAB* and *ΔslkAB ΔpilQ* derivatives. The resulting strains were grown in LB, serially diluted, spotted onto LB agar lacking or containing 50 μg/mL erythromycin, and photographed. The inactivated T4P gene products in each spot dilution assay, *pilC* for (B), *pilA* or *pilY1* for (C), *pilMNOP* for (D), and *tsaP* for (E), are indicated with dotted lines and fainter color in the cartoons on the right. Minor pilins are not designated and PilBTU is not drawn for simplicity. (D) For testing the effect of alignment complex inactivation, *pilQ* was expressed from an ectopic locus as in Fig. 2B to avoid the polar effect on *pilQ* expression by *pilMNOP* mutation. The LB and LB agar contained 1 mM IPTG to induce *pilQ* expression.

To investigate the relationship between erythromycin sensitivity and the twitching motility defect, we looked for mutations that can affect twitching motility by resequencing the genomes of the two PAO1 strains using the PAO1 reference genome (NC_002516) (Stover *et al*, 2000). The two genomes showed differences at a few genetic loci, including a change at the *pilC* locus (Table S1-S2). PilC is the platform protein that coordinates the polymerization and depolymerization of T4P (Takhar *et al*, 2013). Thus, the mutation near *pilC* seemed responsible for the twitching motility defect. After aligning *pilC* coding sequences of PAO1, PAO1^tw^, PA14, and PAK, we realized that the *pilC* start codon was misannotated in the PAO1 reference genome and that there is actually a frameshift mutation in *pilC* of the reference genome and our twitching-defective PAO1 strain by the insertion of four bases (ACTG) after the 27^th^ codon (Fig. S4). Accordingly, ectopic expression of the *pilC* of the PAO1^tw^ strain restored the twitching motility in the twitching-defective PAO1 strain and its *ΔslkAB* derivative (Fig. S5A).

Next, we examined the effect of *pilC* expression on the erythromycin susceptibility of the *ΔslkAB* derivative of the twitching-defective PAO1 strain. To our amazement, *pilC* expression completely suppressed the erythromycin susceptibility, suggesting that the *pilC* mutation is indeed responsible for the difference in the erythromycin susceptibility phenotype between strains (Fig. S5B). To further examine the requirements for antibiotic susceptibility, we tested the effect of combining *pilC* and *slkAB* mutations in the twitching-proficient PAO1^tw^, PA14, and PAK strains. As expected, the *ΔpilC ΔslkAB* strains became susceptible to erythromycin (Fig. 3B and Fig. S3). Moreover, the erythromycin sensitivity of the *ΔpilC ΔslkAB* strains was suppressed by a *pilQ* mutation. These results suggested that the PilQ secretin channel functions as a pore through which antibiotics are translocated when Slk proteins are inactivated and type IV pili assembly is not completed because of a PilC defect.

We next sought to determine if other mutations that cause a defect in T4P assembly also result in antibiotic permeation through the PilQ secretin in the *ΔslkAB* strain. The T4P assembly system is composed of four subcomplexes: the platform protein PilC and cytoplasmic motor proteins PilBTU; the pilus shaft consisting of the major pilin PilA, minor pilins FimU-PilVWXE, and the adhesin PilY1; the alignment complex made up of PilMNOP; and the secretin complex consisting of PilQ, PilF, and TsaP (Burrows, 2012; McCallum *et al*, 2019). We first tested if the lack of the pilus shaft also causes antibiotic permeation in the absence of Slk proteins similar to a *pilC* mutation. Assembly of the pilus shaft begins with formation of a priming complex consisting of an adhesin and minor pilins (Treuner-Lange *et al*, 2020), and the pilus elongates by addition of major pilins at the base of the pilus in the IM (Jacobsen *et al*, 2020). We hypothesized that the loss of the pilus shaft resulting from deletion of the major pilin *pilA* or the adhesin *pilY1* would leave the PilQ secretin pore unoccupied and cause antibiotic permeation through the secretin pore if the Slk proteins are not present. Accordingly, introduction of *pilA* or *pilY1* mutation also caused erythromycin sensitivity in the *ΔslkAB* derivative of the PAO1^tw^ strain, and the sensitivity was suppressed by *pilQ* deletion as observed for *pilC* mutants (Fig. 3C).

To test the effect of mutations in the alignment complex genes on antibiotic permeation through the PilQ secretin, we used a *ΔpilMNOPQ* mutant strain that ectopically expresses *pilQ* from an inducible promoter. This configuration was required because mutations in the *pilMNOP* genes appeared to be polar on the expression of downstream *pilQ* (Ayers *et al*, 2009) (Fig. 2A). The *ΔslkAB ΔpilMNOPQ* mutant strain became susceptible to erythromycin treatment upon induction of *pilQ* expression from an ectopic locus in the chromosome, but showed erythromycin resistance without *pilQ* induction (Fig. 3D). Thus, mutations in genes encoding components of the T4P subcomplexes assembling at the IM seemed to generally cause a PilQ-mediated defect in the OM permeation barrier upon inactivation of the Slk proteins.

### Loss of TsaP also causes PilQ-mediated antibiotic diffusion in the absence of Slk proteins

In contrast to the mutations of the IM subcomplexes, mutations of the genes required for assembly of the OM secretin channel such as *pilF* and *fimV* were identified as suppressors of the erythromycin susceptibility along with *pilQ* mutations (Fig. 2A). TsaP is a component of the T4P secretin complex that forms peripheral spikes surrounding the complex (Siewering *et al*, 2014; Chang *et al*, 2016; McCallum *et al*, 2021). It was shown to be important for T4P surface assembly and twitching motility in *Neisseria gonorrhoeae* and *Myxococcus xanthus* (Siewering *et al*, 2014). However, the PilQ secretin complexes are still formed in the *ΔtsaP* strain irrespective of the twitching motility defect (Siewering *et al*, 2014; Chang *et al*, 2016). Moreover, in *P. aeruginosa*, a *tsaP* mutation did not cause a twitching motility defect as severe as in *N. gonorrhoeae* or *M. xanthus* (McCallum *et al*, 2021). Thus, we suspected that the phenotype related to antibiotic permeation might be different between the *ΔtsaP* strain and other mutant strains of the T4P secretin complex. Indeed, inactivation of TsaP did not suppress the erythromycin susceptibility of the *ΔpilC ΔslkAB* strain (Fig. S6).

TsaP was proposed to function in anchoring the OM secretin complex to the peptidoglycan layer and/or aligning the secretin complex with the IM complex (Siewering *et al*, 2014). We reasoned that the alignment problem would cause a significant fraction of PilQ secretin pores to remain unoccupied in the *ΔtsaP* mutant. If so, a *tsaP* mutation might also cause an OM permeability defect in the absence of Slk proteins. Accordingly, the *ΔtsaP ΔslkAB* mutant became susceptible to erythromycin, and the erythromycin sensitivity was suppressed by *pilQ* mutation (Fig. 3E). An alternative explanation for the permeability defect of the *ΔtsaP ΔslkAB* strain seemed to be that TsaP and the Slk proteins function redundantly for preventing permeation through the T4P secretin complex. However, unlike the *ΔslkAB ΔpilC* double mutant, the *ΔtsaP ΔpilC* double mutant did not exhibit a noticeable increase in erythromycin susceptibility (Fig. S7), suggesting that TsaP does not have a major role in preventing permeation through the PilQ secretin channel but is instead important to prevent the formation of disengaged pores.

### Slk proteins interact with the PilQ secretin complex

The above results led us to hypothesize that the Slk proteins prevent antibiotic permeation by interacting with the PilQ secretin channel when the channel is not docked with the IM complex. As PilQ localizes to the poles independently of the Slk proteins or the IM complex assembly (Fig. S8) (Carter *et al*, 2017), we reasoned that the interaction between Slk proteins and the PilQ channel can be assessed by examining the polar localization of Slk proteins in the PAO1^tw^ strain and its T4P mutant derivatives. To monitor SlkA localization, we used a SlkA-mScarlet fusion protein produced from its native chromosomal locus. As SlkB might interfere with SlkA-mScarlet localization by competing for interaction with the PilQ channel, we deleted *slkB* when we introduced *slkA-mScarlet* fusion at the native locus. As for SlkB localization, *slkB-mScarlet* was expressed from an ectopic locus in the chromosome (the Tn7 integration site) in the *ΔslkAB* strain background to avoid competition between SlkB-mScarlet and untagged Slk proteins. SlkA-mScarlet and SlkB-mScarlet are partially functional in that the strains expressing the fusion proteins do not show a permeability defect when combined with *ΔpilC* that is as severe as the *ΔpilC ΔslkAB* strain (Fig. S9).

In accordance with our hypothesis, SlkA-mScarlet and SlkB-mScarlet localized to the poles in a PilQ-dependent fashion, albeit weakly, indicating that Slk proteins interact with the PilQ secretin complex (Fig. 4 and Fig. S10). Strikingly, the polar localization of Slk proteins was dramatically enhanced when assembly of the IM complex was impaired by *ΔpilC*, suggesting that Slk proteins interact with the PilQ secretin channel more strongly when the IM complex assembly is inhibited. These results imply that the Slk proteins and the IM complex compete for interaction with the PilQ secretin channel and that the Slk proteins are displaced when the IM complex assembles with the secretin channel. Overall, the localization pattern of Slk proteins in wild-type and T4P mutant strains is consistent with our hypothesis that Slk proteins interact with the PilQ secretin channel to prevent the diffusion through the OM when the IM complex is not properly assembled with the secretin channel.

**Figure 4.**
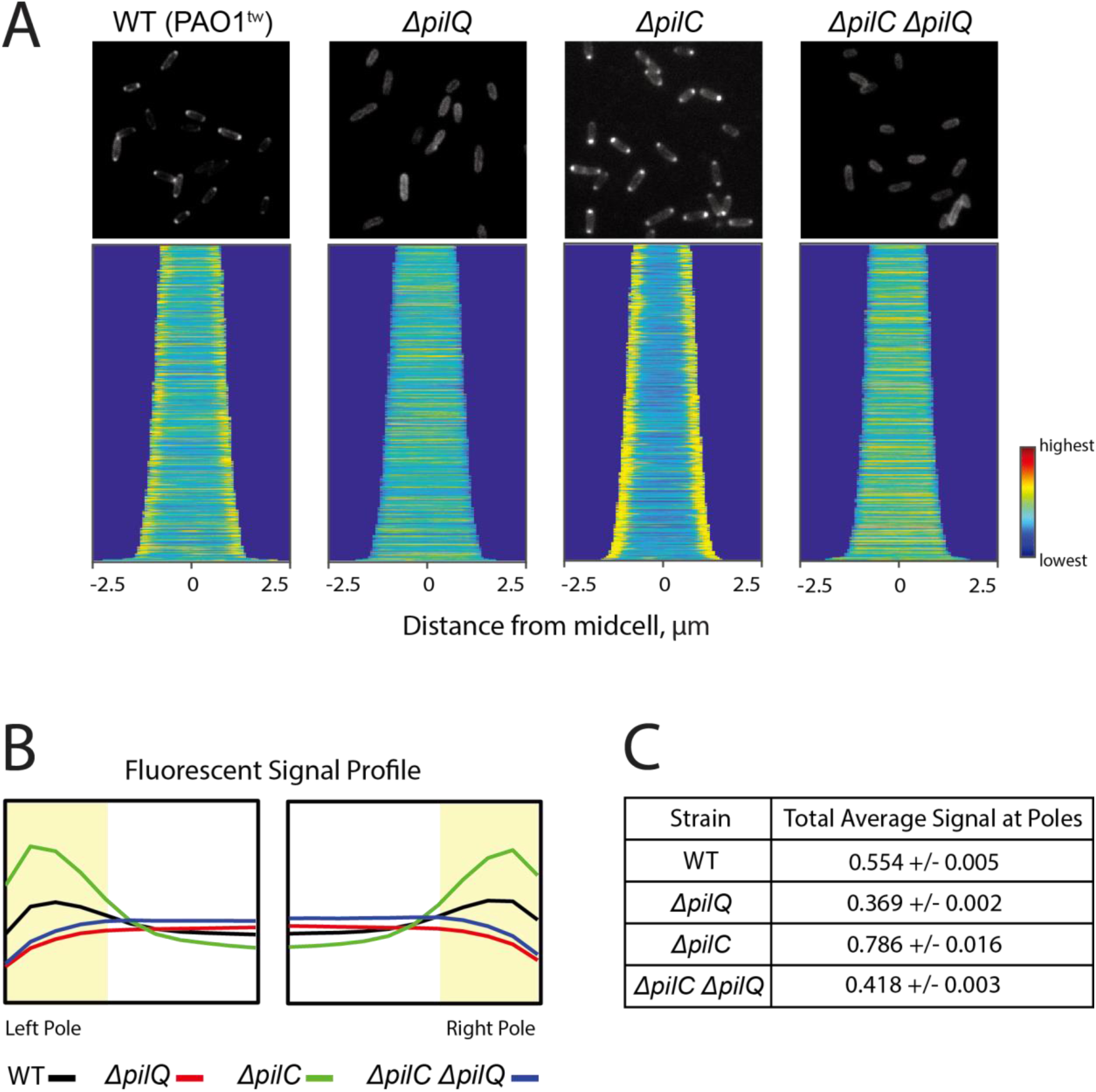
PilQ-dependent polar localization of SlkA and its enhancement upon inactivation of the T4P platform PilC. (A) A PAO1^tw^ derivative that express *slkA*-*mScarlet* from its native locus (OKP59) and its *ΔpilQ*, *ΔpilC*, and *ΔpilC ΔpilQ* derivatives (OKP60, OKP61, and OKP120) were grown overnight in LB. The overnight cultures were diluted 1:100 in M9-glucose (0.2%) medium, grown to exponential phase (OD600 = 0.2∼0.3), and imaged on 2% agarose pads containing 1X M9 salts. Shown on top of each panel are representative fluorescent images showing the localization of SlkA-mScarlet in each strain. Below the micrographs are demographs that reflect SlkA-mScarlet localization throughout a population of 500 cells arranged according to cell length. Single-cell fluorescence quantification was performed using Oufti (59). (B) The mean SlkA-mScarlet signal intensity profile per unit length was calculated for the 500 single-cells of each strain shown in (A). Each plot displays the mean signal every 0.0648 microns for 11 points, spanning from the outermost cell tips inward. The cell pole is defined by the yellow highlighted region corresponding to the outermost 0.324 microns. (C) Quantification of SlkA-mScarlet signal at the poles. Shown are mean and standard deviation of the SlkA-mScarlet fluorescent signal intensity at the cell poles, defined in (B), for each of the indicated strains.

### Slk proteins are likely to function specifically for the T4PS in P. aeruginosa

In *P. aeruginosa*, there are three more virulence systems that assemble OM channels comprised of secretin homologues: two T2SS (Xcp and Hxc) and a T3SS (Psc). As the secretin complexes of these systems are likely to be assembled independently of the subcomplexes in the IM (Diepold & Wagner, 2014; Thomassin *et al*, 2017; Douzi *et al*, 2012), we examined if the Slk proteins are also involved in preventing drug permeation through these secretin channels. The platform protein gene of each system, *xcpS*, *hxcS*, or *pscJ*, was deleted to inhibit the assembly of the IM subcomplexes of the secretin-containing systems in the *ΔslkAB* strain of the PAO1^tw^ strain. The antibiotic sensitivity of the resulting strains was then compared with that of the *ΔslkAB* strain in growth conditions that support the production and assembly of the secretin complexes of each virulence system. For assessing the antibiotic susceptibility of the *ΔhxcS ΔslkAB* strain, a phosphate-limiting proteose peptone medium was used to induce the expression of Hxc T2SS (Ball *et al*, 2002). The effect of a *pscJ* mutation was examined in the strains that overproduce the master regulator ExsA to induce T3SS expression. However, we did not observe increased sensitivity of these strains to macrolides compared with that of the PAO1^tw^ and its *ΔslkAB* derivative (Fig. S11), which suggested that SlkA and SlkB are likely to function specifically with the T4PS.

## Discussion

The cell envelope of Gram-negative pathogens functions as an effective permeability barrier against antibiotics because the OM efficiently restricts the influx of various toxic compounds and efflux pumps expel the toxic molecules from the periplasm as well as from the cytoplasm. Nevertheless, diffusion of antibiotics across the OM occurs inevitably because Gram-negative bacteria assemble channel structures in the OM to carry out various cellular functions important for survival and virulence. General porins and substrate-specific channels required for nutrient transport have been implicated as a major route for diffusion of small and hydrophilic drugs such as beta-lactams (Vergalli *et al*, 2020; Zgurskaya *et al*, 2015). Secretin complexes assembled in the OM as part of protein secretion machineries are another type of channel structures that can potentially serve as a conduit for drug diffusion, but mechanistic details related to the permeation of compounds through these structures have remained unclear. In this study, we found that defects of the IM complex assembly or alignment between the IM and OM complexes of *P. aeruginosa* T4P assembly systems can cause diffusion of drugs through the OM secretin channel. We also discovered that in these situations, drug diffusion through the secretin channel is prevented by periplasmic proteins SlkA and SlkB that interact specifically with the channel complex until the T4PS is fully assembled.

Although secretin channels form large pores to accommodate protein substrates, diffusion through the channels is thought to be minimized by the gate structure of the channels that are closed unless proteins are secreted through the channel. However, our results indicate that the gate structure does not effectively seal the T4P secretin channel against certain classes of antibiotics such as macrolides and aminoglycosides in *P. aeruginosa* when the IM complex is not docked to the channel. It has been assumed that there are “plug” proteins that help seal secretin channels more tightly when the channels are not engaged in their function. Supporting this idea, electron densities for plug-like proteins have been visualized in several EM studies of secretin complexes, but the identity of the plug proteins has not been determined yet (Chami *et al*, 2005; Chernyatina & Low, 2019; Ghosal *et al*, 2019; McCallum *et al*, 2021). A recent cryo-electron tomography analysis of T4P in *Myxococcus xanthus* suggested that PilY1 is likely to function as a plug at the entry to the PilQ secretin vestibule based on the tomograms taken in the cells lacking specific domains of T4P components (Treuner-Lange *et al*, 2020). Thus, after the assembly of T4PS, the proteins that are located at the top of the pilus shaft structure such as PilY1 appear to function as a plug that seals the secretin channel in the retracted pili, while the channel is likely to be sealed by the pilus shaft itself in the extended pili (Fig. 5). However, the plug structure was only visible in the fully assembled T4PS, suggesting that the plug is installed when the IM complex assembles with the secretin channel. Thus, it remained unclear whether there are plug proteins that seal the secretin channel when the transenvelope complex is not fully assembled.

**Figure 5.**
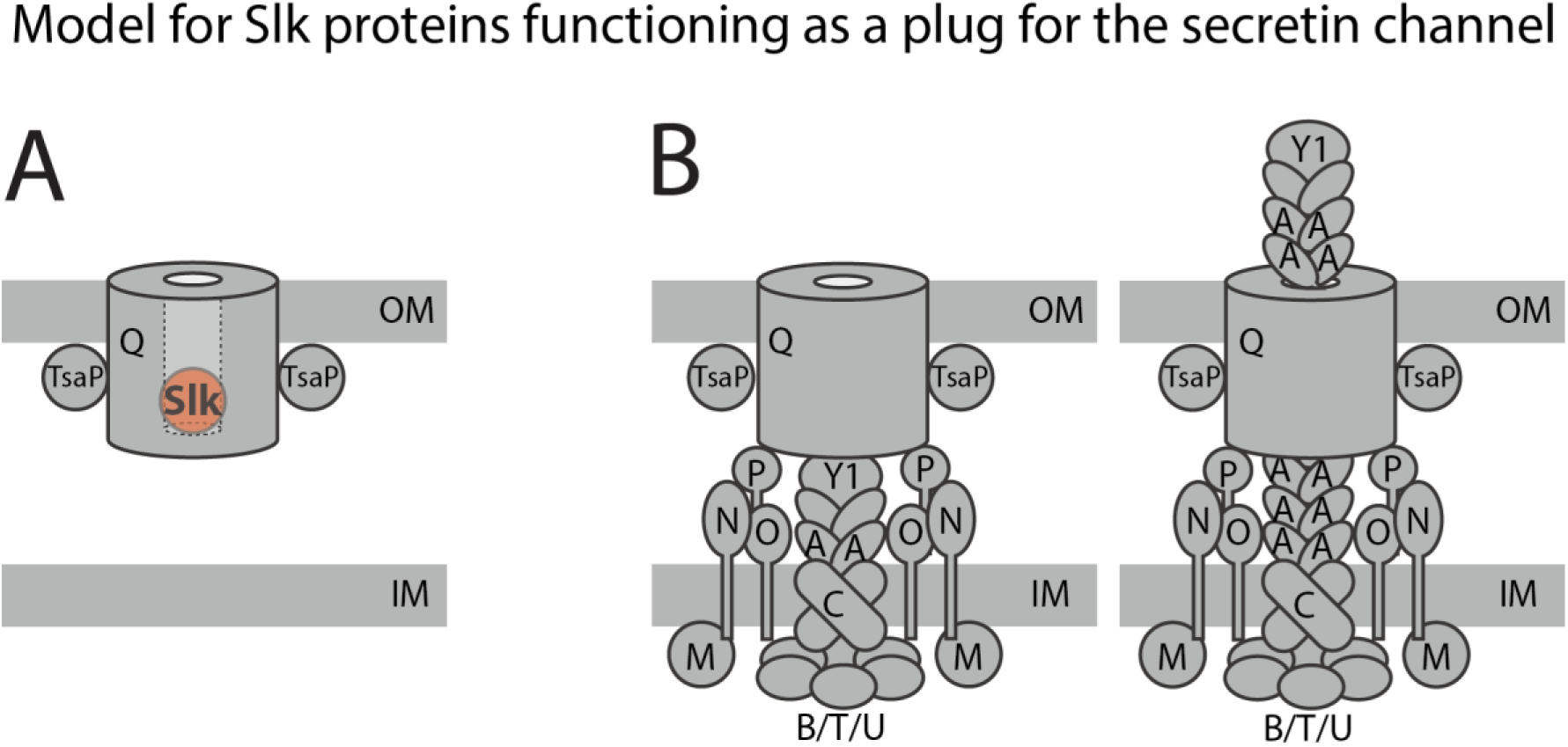
Model for Slk proteins functioning as a plug that prevent leakage through the OM secretin complex of the T4P. (A) The large, cylindrical figure represents the multimeric OM channel consisting of PilQ secretin and TsaP proteins. The gate structure of the secretin channel is omitted for simplicity. Slk proteins indicated as a red circle prevent diffusion of compounds through the channel until the IM complex docks with the secretin channel. (B) Slk proteins are displaced when the T4P is fully assembled. After the full assembly of T4P, some components of the IM complexes are likely to function as a plug in the complexes with retracted pili, while the pili itself seals the channel in the complexes with extended pili.

Results from our genetic and microscopic analyses indicate that the Slk proteins function as a plug for the secretin channels that are not docked with the IM complexes. The *slkAB* mutant showed an OM barrier defect, but the defect only became obvious when the IM complex could not be assembled or aligned with the OM secretin channel. This result suggested that either the Slk proteins or the IM complex can prevent the diffusion through the secretin channel, and thus the activity of the Slk proteins is not critical when the T4PS is efficiently assembled. In addition, when we assessed the interaction of Slk proteins with the secretin complex using the PilQ-dependent polar localization as an interaction proxy, inhibition of the IM complex assembly strongly enhanced the polar localization of Slk proteins (Fig. 4 and Fig S10). Moreover, Slk proteins were completely delocalized in the *pilQ* mutant background irrespective of the IM complex mutation, indicating that the enhanced polar localization is also dependent on PilQ. Thus, while it is formally possible that Slk proteins are recruited to the poles via interactions with other proteins associated with the PilQ secretin channel, these results strongly suggested that they engage with the secretin channel before the IM complex docks to the channel and are displaced from the channel upon full T4PS assembly. SlkA (PA5122) was also serendipitously detected in a sample of *P. aeruginosa* PilQ secretin complex prepared for cryo-EM (McCallum *et al*, 2021). Although the role of SlkA was not appreciated by the authors at the time, their result provides independent support for the specific interaction between SlkA and the PilQ secretin channel. Taken together, all the available data are consistent with the idea that the Slk proteins function as a plug that help seal the secretin channel to maintain the OM permeability barrier until the T4PS is fully assembled (Fig. 5).

SlkAB homologues are only found in gamma-proteobacteria and not widely conserved among bacterial species that have the T4PS or other secretin-containing virulence systems (Fig. S12-13), but we speculate that different types of plug proteins for secretin channels are likely to exist among bacteria that lack SlkAB homologues. In many secretin-reliant systems, assembly of secretin complexes occurs independently of the IM complex assembly (Korotkov *et al*, 2011; Diepold & Wagner, 2014; Carter *et al*, 2017; Friedrich *et al*, 2014; Douzi *et al*, 2012; Thomassin *et al*, 2017). Thus, defective assembly of the IM complex or improper alignment between the OM and IM complexes in the secretin-dependent systems might also cause a breach in the permeability barrier function of the OM. Proteins that help seal the secretin pores are therefore also likely to exist among other secretin-dependent virulence systems. In this regard, it is noteworthy that other important secretin interacting proteins are also divergent in sequence. For example, the pilotins that deliver secretins to the OM and/or aid in secretin assembly belong to several groups of proteins that are unrelated in their sequence and structure even though they serve a similar function (Silva *et al*, 2020; Koo *et al*, 2012). We thus suspect that proteins that prevent diffusion through secretin channels might also be diverse among secretin-dependent systems.

Because we identified the OM permeability defect of the *slk* mutant with macrolides that are lipophilic, we initially suspected that the T4P secretin channel might preferentially facilitate the diffusion of hydrophobic drugs. However, hydrophobicity did not seem to be a general property of the chemicals that permeate through the secretin channel. Instead, this channel increased susceptibility of *P.aeruginosa* to several classes of drugs with different properties. Thus, although the loss of Slk protein function combined with defects in T4P assembly clearly creates a PilQ-dependent OM permeability defect, the precise chemical properties that promote permeation through the secretin channel remain to be determined. Understanding these and other chemical properties that promote OM permeation will be critical for the development of future antibiotic therapies effective against Gram-negative pathogens.

OM secretin channels are essential components of many virulence systems and thus have been considered as attractive targets for development of antivirulence drugs (Baron, 2010; Korotkov *et al*, 2011). Our discovery that the PilQ channel functions as a pore for drug diffusion in the *slkAB* mutant indicates that secretin channels can also be exploited as a conduit that facilitate drug delivery into Gram-negative pathogens. Thus, understanding how secretin-reliant systems prevent diffusion of toxic compounds through secretin channels is a promising avenue for developing effective strategies that broaden the spectrum of several approved antibiotics to include activity against Gram-negative infections.

## Materials and Methods

### Media, Bacterial Strains, and Plasmids

Cells were grown in LB (1% tryptone, 0.5% yeast extract, 0.5% NaCl), VBMM, minimal M9 medium supplemented with 0.2 % glucose and 0.2 % casamino acids, or proteose peptone medium (Cheng *et al*, 1970). The strains and plasmids that were used for the study are listed in *SI Appendix*, Table S3 and Table S4. Detailed procedures for strain and plasmid construction are also provided in *SI Appendix*.

### Transposon sequencing

*P. aeruginosa* PAO1 was mutagenized with the mariner transposon from pTBK30 (Goodman *et al*, 2004). Genomic DNA from each cell pellet was extracted, fragmented, poly-C tailed, and sequenced as previously described (Lai *et al*, 2017). Detailed procedure is provided in *SI Appendix*.

### Antibiotic susceptibility testing – spot dilution assay

Overnight grown strains of interests were diluted to OD_600_=2.0, serially diluted 10-fold, and 5 μl of each dilution was spotted on LB agar supplemented with indicated antibiotics. The plates were photographed after incubation for 20 hours at 37 °C, unless otherwise specified.

### Antibiotic susceptibility testing – agar diffusion assay

Freshly saturated cultures were diluted to OD_600_ = 2.0, and 125 μL of the normalized cultures was mixed with 5 mL molten H-top soft agar (1% tryptone, 0.8% NaCl, 0.7% agar) and spread on 10 centimeter diameter LB agar plates. Then, MIC Test Strips (Liofilchem) were applied to the solidified soft agar surface. Alternatively, antibiotics were serially diluted and 5 μL of each dilution was spotted on the soft agar surface. The plates were incubated for 24 hours at 37 °C before being photographed.

### Microscopic image acquisition and analysis

Growth conditions prior to microscopy are described in the figure legends. Prior to imaging, cells were immobilized on 2% agarose pads containing 1X M9 salts and covered with #1.5 coverslips. Micrographs were obtained using a Leica DM2500 LED microscope equipped with a Leica DFC7000 GT camera, Fluo Illuminator LRF 4/22, HC PL APO 100x/1.40 Oil Ph3 objective lens, and Leica Las X acquisition software. Images in the red channel were obtained using N2.1 filter cube. Automated cell segmentation and identification as well as measurements of fluorescence signal at the single cell level were carried out using Oufti (Paintdakhi *et al*, 2016). For demographs, custom-written MATLAB code was used to arrange cells from top to bottom according to their length as previously described (Sher *et al*, 2020).

## ACKNOWLEDGMENTS.

We thank the members of Cho lab at Sungkyunkwan University for helpful comments and discussion. This study was supported by the National Research Foundation of KOREA (NRF-2019R1A2C1002648) and the Basic Science Research Program of the Ministry of Education (NRF-2019R1A6A1A10073079).

## SUPPLEMENTAL MATERIALS AND METHODS

### Media, Bacterial Strains, and Plasmids

Cells were grown in either LB [1% tryptone, 0.5% yeast extract, 0.5% NaCl], Vogel-Bonner minimal medium (VBMM) [3.42 g/L trisodium citrate dihydrate, 2 g/L citric acid, 10 g/L K2HPO4, 3.5 g/L NaNHPO4-4H2O, pH7, 1 mM MgSO4, 0.1 mM CaCl2], minimal M9 medium supplemented with glucose [6 g/L Na2HPO4 anhydrous, 3 g/L KH2PO4, 0.5 g/L NaCl, 1 g/L NH4Cl, 0.2 % glucose, 2 mM MgSO4, 0.1 mM CaCl2], or proteose peptone medium [20 mM NH4Cl, 20 mM KCl, 120 mM Tris-HCl, pH7.4, 0.5% glucose, 0.5% proteose peptone (Difco), and 1.6 mM MgSO4] (Cheng *et al*, 1970). The strains and plasmids that were used for the study are listed in Table S3 and Table S4. Detailed procedures for strain and plasmid construction are also provided in the Supporting Information.

### Plasmid construction

#### SpBO37

To make a vector for expression of periplasmic C-terminal mScarlet fusion proteins, an 838 bp g-block DNA fragment containing φ10 RBS-NcoI site-the signal sequence of PAO1 *dsbA*-BamHI site-PstI site-5 amino acid linker-mScarlet coding sequence between BglII and KpnI sites was synthesized from IDT. The g-Block DNA was digested with BglII and KpnI and ligated with pKHT4 digested with BamHI and KpnI, which make the BamHI site between dsbA signal sequence and mScarlet unique for later use.

#### pBO48

To clone *exsA* under arabinose promoter, *exsA* gene with native RBS was amplified using a primer pair 5’-GCTA*<u>GAATTC</u>*GTACGACGGGAAGTGTTGG -3’ and 5’-GCTA*<u>TCTAGA</u>*CGTCAGTTATTTTTAGCCCGG-3’. The resulting PCR product was digested with EcoRI and XbaI and ligated with pJN105 digested with the same enzymes to generate pBO048.

#### pOKP10

To construct a plasmid for deletion of *slkA* (*PA5122*), a 700bp region upstream of *slkA* was amplified using 5’-GCTAAAGCTTCTGGGAAAGCGCTCCTCGAG-3’ and 5’-TTTCTTCCCGAGGTGCATGGACGGTCTCCGGGTGA-3’. A 701 bp region downstream of *slkA* was amplified using 5’-CGGAGACCGTCCATGCACCTCGGGAAGAAACGATG-3’ and 5’-GCTATCTAGAGGTAATCGCACCAATCTGGTG-3’. The resulting PCR products were spliced using overlap extension PCR (OE) using the primers 5’-GCTAAAGCTTCTGGGAAAGCGCTCCTCGAG-3’ and 5’-GCTATCTAGAGGTAATCGCACCAATCTGGTG-3’. The OE product was digested with HindIII and XbaI and ligated with pEXG2 digested with the same enzymes to generate pOKP10.

#### pOKP11

To construct a plasmid for deletion of *slkB* (*PA5123*), a 702bp region upstream of *slkB* was amplified using 5’-GCTAAAGCTTCTGGCGCAACGAGGGTTC-3’ and 5’-CTCGCTTGGCTCTATCATCGTTTCTTCCCGAGGTG-3’. A 700 bp region downstream of *slkB* was amplified using 5’-CGGGAAGAAACGATGATAGAGCCAAGCGAGCGCTA-3’ and 5’-GCTATCTAGAGCACCAGCAGCTTGGTGGTC-3’. The PCR products were spliced by OE using the primers 5’-GCTAAAGCTTCTGGCGCAACGAGGGTTC-3’ and 5’-GCTATCTAGAGCACCAGCAGCTTGGTGGTC-3’. The OE product was cloned between HindIII and XbaI sites of pEXG2 as described above to generate pOKP11.

#### pOKP12

To construct a plasmid for deletion of *slkAB* (*PA5122-PA5123*), a 700bp region upstream of *slkA* was amplified using 5’-GCTAAAGCTTCTGGGAAAGCGCTCCTCGAG-3’ and 5’-CTCGCTTGGCTCTATCATGGACGGTCTCCGGGTG-3’. A 700 bp region downstream of *slkB* was amplified using 5’-CGGAGACCGTCCATGATAGAGCCAAGCGAGCGCTAG-3’ and 5’-GCTATCTAGAGCACCAGCAGCTTGGTGGTC-3’. The PCR products were spliced by OE using the primers 5’-GCTAAAGCTTCTGGGAAAGCGCTCCTCGAG-3’ and 5’-GCTATCTAGAGCACCAGCAGCTTGGTGGTC-3’. The OE product was cloned between HindIII and XbaI sites of pEXG2 as described above to generate pOKP12.

#### pOKP17-pOKP18

To clone *slkA* under arabinose promoter, *slkA* gene with native RBS was amplified using a primer pairs 5’-GTCAGAATTCGGTTCACCCGGAGACCGTCC-3’ and 5’-GTCATCTAGATCATCGTTTCTTCCCGAGGTGC-3’. For *slkB* cloning, *slkB* with native RBS was amplified with 5’-GTCAGAATTCGGTTCACCCGGAGACCGTCCATGATCCCGCGCATTCTCG-3’ and 5’-GTCATCTAGACTAGCGCTCGCTTGGCTC-3’. Each PCR product was digested with EcoRI and XbaI and ligated with pJN105 digested with the same enzymes to generate pOKP17 (*slkA*) and pOKP18 (*slkB*).

#### pOKP23

To construct a plasmid for a *pilQ* deletion, a 620 bp DNA upstream of *pilQ* was amplified using 5’-GCTAAGCTTGGCCAAGACCTATCGCTAC-3’ and 5’-CACCTTGACTTCGACCATCGTCCGACTCCGTTGTA-3’. A 620 bp region including a 300 bp *pilQ* sequence and a 320 bp region downstream of pilQ was amplified using 5’-CGGAGTCGGACGATGGTCGAAGTCAAGGTGACCAA-3’ and 5’-GCTATCTAGACGAGGCATGCAGGTACAC-3’. PCR products were spliced by OE using the primers 5’-GCTAAGCTTGGCCAAGACCTATCGCTAC-3’ and 5’-GCTATCTAGACGAGGCATGCAGGTACAC-3’. The OE product was cloned into pEXG2 using HindIII and XbaI sites to generate pOKP12.

#### pOKP26

To clone GS linker-*mScarlet* under arabinose promoter, *mScarlet* gene with a GS linker was amplified with a primer pair 5’-GTCA*<u>GGATCC</u>*GGTAGCGTTTCTAAAGGTGAAGCAGTTATC-3’ and 5’-GTCA*<u>TCTAGA</u>*TTACTTATACAGTTCATCCATACCTC-3’ using pBO37 as a template. The resulting PCR product was digested with BamHI and XbaI and ligated with pOKP16 digested with the same enzymes to generate pOKP26.

#### pOKP28-pOKP29

A PilQ-mScarlet sandwich fusion was constructed as described in Carter et al (2017) with mScarlet located between the first and second AMIM domains of PilQ. The 416 bp region including 20 bp upstream of *pilQ* and 396 bp *pilQ* coding sequence was amplified using a primer pair 5’-GTCA*<u>GAATTC</u>*CTGTACAACGGAGTCGGACG-3’ and 5’-TTCACCTTTAGAAACGCTACCGACGCTGGCGCCGGCC-3’. *mScarlet* sequence was amplified with 5’-GCCAGCGTCGGTAGCGTTTCTAAAGGTGAAGCAGTTATC-3’ and 5’-GTCA*<u>GGATCC</u>*CTTATACAGTTCATCCATACC-3’ using pBO37 as a template. The PCR products were spliced by OE with a primer pair 5’-GTCA*<u>GAATTC</u>*CTGTACAACGGAGTCGGACG-3’ and 5’-GTCA*<u>GGATCC</u>*CTTATACAGTTCATCCATACC-3’. The overlap extension product was cloned into pOKP16 using EcoRI and BamHI sites to generate pOKP28 that express a truncated *pilQ-mScarlet* fusion gene. The 1,749 bp coding sequence of *pilQ* at the 3’ end was amplified with a primer pair 5’-GTCA*<u>GGATCC</u>*GCTTCCGCCGCTCCGGTC-3’ and 5’-GTCA*<u>TCTAGA</u>*TCAGCGACCGATTGCGATGG-3’. The PCR product was digested with BamHI and XbaI and ligated with pOKP28 digested with the same enzymes to generate pOKP29 that expresses *pilQ-mScarlet* sandwich fusion under arabinose promoter.

#### pOKP35

To construct a plasmid for generating *pilQ-mScarlet* sandwich fusion at the native *pilQ* locus, a 1552 bp region of *pilQ*-*mScarlet* sandwich fusion was amplified with a primer pair 5’-GCT*<u>AAGCTT</u>*CTGTACAACGGAGTCGGACG-3’ and 5’-GCTA*<u>TCTAGA</u>*TTGTCCTTCTTGCGCCGCTC-3’ using pOKP29 as the template. The PCR product was cloned into pEXG2 using HindIII and XbaI sites to generate pOKP35.

#### pOKP42

To construct a plasmid for a *pilC* deletion, a 707 bp sequence upstream of *pilC* from the *pilC* start codon was amplified using 5’-GCTA*<u>TCTAGA</u>*TATCGCTATACACCGGCCTG -3’ and 5’-ATCGAGCGAACCGGACATGGATTAATCCTTGGTCAC-3’. A downstream 717 bp region, including 186 bp *pilC* coding sequence at the 3’ end was amplified using 5’-AAGGATTAATCCATGTCCGGTTCGCTCGATGAGATG -3’ and 5’-GCTA*<u>GGATCC</u>*GAAGAGGCCGAAATGGTTGG-3’. The PCR products were spliced by OE using the primers 5’-GCTA*<u>TCTAGA</u>*TATCGCTATACACCGGCCTG -3’ and 5’-GCTA*<u>GGATCC</u>*GAAGAGGCCGAAATGGTTGG -3’. The OE product was cloned into pEXG2 using BamHI and XbaI.

#### pOKP82

To construct a plasmid for a *pilA* deletion, a 689 bp region upstream of *pilA* including the 49 bp *pilA* sequence at the 5’ end was amplified using a primer pair 5’-GCTA*<u>AAGCTT</u>*CAAGCTTGTCGTCTTCGACC-3’ and 5’-GCAGTACGGTTCAGACAACCACGATCATCAGTTCG-3’. A 647 bp region downstream of *pilA* including 82 bp *pilA* coding sequence at the 3’ end was amplified with a primer pair 5’-TGATGATCGTGGTTGTCTGAACCGTACTGCGGATG-3’ and 5’-GCTA*<u>TCTAGA</u>*CCTCTACGATGCCTTCCTG-3’. The PCR products were spliced by OEusing a primer pair 5’-GCTA*<u>AAGCTT</u>*CAAGCTTGTCGTCTTCGACC-3’ and 5’-GCTA*<u>TCTAGA</u>*CCTCTACGATGCCTTCCTG-3’. The resulting OE product was cloned into pEXG2 using BamHI and XbaI.

#### pOKP88

To construct a plasmid for deletion of *pilMNOPQ* operon, a 620 bp upstream region from *pilM* start codon was amplified using a primer pair 5’-GCTAAAGCTTCTTTCGCGAGAAGCTTCTTTC-3’ and 5’-ACCTTGACTTCGACCACGACCAATTCCCTATTAGC-3’. A 620 bp downstream region including 300 bp *pilQ* coding sequence at the 3’ end was amplified using 5’-AGGGAATTGGTCGTGGTCGAAGTCAAGGTGACCAA-3’ and 5’-GCTATCTAGACGAGGCATGCAGGTACAC-3’. PCR products were spliced by OE using a primer pair 5’-GCTAAAGCTTCTTTCGCGAGAAGCTTCTTTC -3’ and 5’-GCTATCTAGACGAGGCATGCAGGTACAC-3’. The resulting OE product was clone into pEXG2 using BamHI and XbaI.

#### pOKP92

To construct a plasmid for a *slkAB* (*PA14_RS27590-PA14_RS27595*) deletion in PA14, a 700 bp upstream region from the start codon of *PA14_RS27590* (*slkA*) was amplified using 5’-GCTAAAGCTTCTGGGAAAGCGCTCCTCGAG-3’ and 5’-GCTAGCGCTCGCTCATGGACGGTCTCCGGGTG-3’. A 691 bp downstream region including 12 bp *PA14_RS27595* (*slkB*)coding sequence at the 3’ end was amplified using 5’-GGAGACCGTCCATGAGCGAGCGCTAGCCGAAC-3’ and 5’-GCTATCTAGAGCACCAGCAGCTTGGTGGTC-3’. The PCR products were spliced by a primer pair 5’-GCTAAAGCTTCTGGGAAAGCGCTCCTCGAG-3’ and 5’-GCTATCTAGAGCACCAGCAGCTTGGTGGTC-3’. The resulting OE product was clone into pEXG2 using BamHI and XbaI.

#### pOKP93

To construct a plasmid for a *slkAB* (*Y880_RS24175-Y880_RS24180*) deletion in PAK, a 699 bp upstream region from the start codon of *Y880_RS24175* (*slkA*) was amplified using 5’-GCTAAAGCTTTTGGAAAGCGCTCCTCGAG-3’ and 5’-CTCGCTTGGCTCTATCATGGACGGTCTCCGGGTG-3’. A 700 bp downstream region including 21 bp *Y880_RS24180* (*slkB*) coding sequence at the 3’ end was amplified using 5’-CGGAGACCGTCCATGATAGAGCCAAGCGAGCGCTAG-3’ and 5’-GCTATCTAGAGCACCAGCAGCTTGGTGGTC-3’. The PCR products were spliced by OE using a primer pair 5’-GCTAAAGCTTTTGGAAAGCGCTCCTCGAG-3’ and 5’-GCTATCTAGAGCACCAGCAGCTTGGTGGTC-3’. The resulting OE product was clone into pEXG2 using BamHI and XbaI.

#### pOKP94

o construct a plasmid for a *pilC* deletion in PA14, a 234 bp region upstream of *PA14_ RS23960* (*pilC*) was amplified using 5’-GCTA*<u>AAGCTT</u>*GTGGGCTGCGACCATTGC-3’ and 5’-TTGTCAGGTTGTCGACATGGATTAGTCCTTGGTCAC-3’. A 306 bp downstream region, including 116 bp *PA14_ RS23960* (*pilC*) sequence was amplified using 5’-AAGGACTAATCCATGTCGACAACCTGACAACGTTG-3’ and 5’-GCTA*<u>TCTAGA</u>*TAGGTCGCCTGCTTGGGTTC-3’. PCR products were spliced by OE using a primer pair 5’-GCTA*<u>AAGCTT</u>*GTGGGCTGCGACCATTGC-3’ and 5’-GCTA*<u>TCTAGA</u>*TAGGTCGCCTGCTTGGGTTC-3’. The OE product was cloned into pEXG2 using HindIII and XbaI sites to generate pOKP94.

#### pOKP95

To construct a plasmid for a *pilC* deletion in PAK, a 615 bp region upstream of *Y880_ RS20950* (*pilC*) was amplified using 5’-GCTA*<u>AAGCTT</u>*AACCAGGTCAACGTCAATCC-3’ and 5’-ATCGGTTCCATGAGCATGGATTAATCCTTGGTCACG-3’. A 633 bp downstream region including a 102 bp *Y880_ RS20950* sequence at the 3’ end was amplified using 5’-AGGATTAATCCATGCTCATGGAACCGATGATCATG-3’ and 5’-GCTA*<u>GGATCC</u>*GAAGAGGCCGAAATGGTTGG-3’. The PCR products were spliced by OE using the primers 5’-GCTA*<u>AAGCTT</u>*AACCAGGTCAACGTCAATCC-3’ and 5’-GCTA*<u>GGATCC</u>*GAAGAGGCCGAAATGGTTGG-3’. The resulting OE product was cloned into pEXG2 using HindIII and BamHI sites to generate pOKP93.

#### pOKP98

To construct a plasmid for a *fimV* deletion, a 750 bp region upstream of *fimV* was amplified using a primer pair 5’-GCTA*<u>AAGCTT</u>*CAACTGAACCTGACAGCCTG-3’ and 5’-TGGCTGTCATTACCTCAGTGTACGAAGCCGAACC-3’. A 719 bp downstream region including a 55 bp *fimV* sequence at the 3’ end was amplified using a primer pair 5’-CGGCTTCGTACACTGAGGTAATGACAGCCAGCAGG-3’ and 5’-GCTA*<u>TCTAGA</u>*TGCGCACCATATGGTGAAGG-3’. PCR products were spliced by OE using the primers 5’-GCTA*<u>AAGCTT</u>*CAACTGAACCTGACAGCCTG-3’ and 5’-GCTA*<u>TCTAGA</u>*TGCGCACCATATGGTGAAGG-3’. The OE product was cloned into pEXG2 using HindIII and XbaI sites to generate pOKP98.

#### pOKP99

To construct a plasmid for a *tsaP* deletion, a 653 bp region upstream of *tsaP* was amplified using 5’-GCTA*<u>AAGCTT</u>*CACGCCTGCTGTCGATG-3’ and 5’-TGGCCATCAGAACCATTTCCTCATGCAGTGAATCC-3’. A 536 bp downstream region, including a 56 bp *tsaP* sequence at the 3’ end was amplified using a primer pair 5’-CACTGCATGAGGAAATGGTTCTGATGGCCAGCAG-3’ and 5’-GCTA*<u>TCTAGA</u>*GTTCCAGAGTCTCCGGAG-3’. The PCR products were spliced by OE using the primers 5’-GCTA*<u>AAGCTT</u>*CACGCCTGCTGTCGATG-3’ and 5’-GCTA*<u>TCTAGA</u>*GTTCCAGAGTCTCCGGAG-3’. The OE product was cloned into pEXG2 using HindIII and XbaI sites to generate pOKP99.

#### pOKP101

To construct a plasmid for generating *slkA-mScarlet* with a *slkB* deletion at the native *slkAB* locus, a 1262 bp *slkA*-*mScarlet* sequence was amplified with a primer pair 5’-GCTA*<u>AAGCTT</u>*GTGCAACGCTTGGCAGGTTC-3’ and 5’-TAGGTGTAGACCCCTTACTTATACAGTTCATCCATACCTC-3’ using pOKP57 a template. A 516 bp region of 5’ end-truncated *slkB* was amplified using 5’-GAACTGTATAAGTAAGGGGTCTACACCTACATCG-3’ and 5’-GCTA*<u>TCTAGA</u>*CTAGCGCTCGCTTGGCTC-3’. PCR products were spliced by overlap extension PCR using the primers 5’-GCTA*<u>AAGCTT</u>*GTGCAACGCTTGGCAGGTTC-3’ and 5’-GCTA*<u>TCTAGA</u>*CTAGCGCTCGCTTGGCTC-3’. The overlap extension product was cloned into pEXG2 using HindIII and XbaI sites to generate pOKP101.

#### pOKP103

To construct a plasmid for an *oprM* deletion, a 700 bp region upstream of *oprM* was amplified using 5’-GCTA*<u>AAGCTT</u>*GAGCGCTACAATGGCGTG-3’ and 5’-GGAACGCCGTCTGGACATATCATTGCCCCTTTTCG-3’. A 628 bp downstream region including a 323 bp *oprM* sequence at the 3’ end was amplified using 5’-AGGGGCAATGATATGTCCAGACGGCGTTCCAGG-3’ and 5’-GCTA*<u>TCTAGA</u>*CAGGGTCGCGAGAATGC-3’. The PCR products were spliced by OE using the primers 5’-GCTA*<u>AAGCTT</u>*GAGCGCTACAATGGCGTG-3’ and 5’-GCTA*<u>TCTAGA</u>*CAGGGTCGCGAGAATGC-3’. The OE product was cloned into pEXG2 using HindIII and XbaI sites to generate pOKP103.

#### pOKP107

To construct a plasmid for a *hxcS* deletion, a 721 bp region upstream of *hxcS* was amplified using 5’-GCTA*<u>AAGCTT</u>*TGGACAACCTGATCCGCCAG-3’ and 5’-GGTGGGTGAGGATCGCATCAGGCGTCCCGGGTC-3’. A 698 bp downstream region including a 221 bp *hxcS* sequence at the 3’ end was amplified with a primer pair 5’-CCGGGACGCCTGATGCGATCCTCACCCACCTGATC-3’ and 5’-GCTA*<u>TCTAGA</u>*GGAAGTCGTCTTGCATGAGG-3’. The PCR products were spliced by OE using a primer pair 5’-GCTA*<u>AAGCTT</u>*TGGACAACCTGATCCGCCAG-3’ and 5’-GCTA*<u>TCTAGA</u>*GGAAGTCGTCTTGCATGAGG-3’. The resulting OE product was cloned into pEXG2 using HindIII and XbaI sites to generate pOKP107.

#### pOKP115

To construct a plasmid for a *pilY1* deletion, a 514 bp region upstream of *pilY1* was amplified using 5’-GCTA*<u>AAGCTT</u>*GCATGCGCGAAGTGGTAC-3’ and 5’-CAGGGGTCATCGTTCCATGCGAGGCTCGATCAG-3’. A 509 bp downstream region including a 313 bp *pilY1* sequence at the 3’ end was amplified using 5’-ATCGAGCCTCGCATGGAACGATGACCCCTGTGC-3’ and 5’-GCTA*<u>TCTAGA</u>*CTTGACCTGGAACAGGCTG-3’. The PCR products were spliced by OE using a primer pair 5’-GCTA*<u>AAGCTT</u>*GCATGCGCGAAGTGGTAC-3’ and 5’-GCTA*<u>TCTAGA</u>*CTTGACCTGGAACAGGCTG-3’. The OE product was cloned into pEXG2 using HindIII and XbaI sites to generate pOKP115

#### pOKP116

To construct a plasmid for a *pilQ* deletion in PA14, a 234 bp region upstream of *PA14_ RS27185* (*pilQ*) was amplified using 5’-GCT*<u>AAGCTT</u>*TTACTTATAGCGCTTCGAACC-3’ and 5’-CCTTCACCTCAACGACATCGTCCGACTCCGTTGTA-3’. A 622 bp downstream region including a 302 bp *PA14_ RS27185* sequence at the 3’ end was amplified using 5’-CGGAGTCGGACGATGTCGTTGAGGTGAAGGTGACC-3’ and 5’-GCTA*<u>TCTAGA</u>*CGAGGCATGCAGGTACAC-3’. The PCR products were spliced by OE using a primer pair 5’-GCT*<u>AAGCTT</u>*TTACTTATAGCGCTTCGAACC-3’ and 5’-GCTA*<u>TCTAGA</u>*TAGGTCGCCTGCTTGGGTTC-3’. The resulting OE product was cloned into pEXG2 using HindIII and XbaI sites to generate pOKP116.

#### pOKP117

To construct a plasmid for a *pilQ* deletion in PAK, a 620 bp region upstream of *Y880_ RS23755* (*pilQ*) was amplified using 5’-GCT*<u>AAGCTT</u>*GGCCAAGACCTATCGCTAC-3’ and 5’-CACCTTGACTTCGACCATCGTCCGACTCCGTTGTA-3’. A 620 bp downstream region, including a 300 bp *Y880_ RS23755* sequence at the 3’ end was amplified using 5’-CGGAGTCGGACGATGGTCGAAGTCAAGGTGACCAA-3’ and 5’-GCTA*<u>TCTAGA</u>*CGAGGCATGCAGGTACAC-3’. The PCR products were spliced by OE using a primer pair 5’-GCT*<u>AAGCTT</u>*GGCCAAGACCTATCGCTAC-3’ and 5’-GCTA*<u>TCTAGA</u>*CGAGGCATGCAGGTACAC-3’. The resulting OE product was cloned into pEXG2 using HindIII and BamHI sites to generate pOKP117.

#### pOKP121

For constructing a plasmid that express *pilQ* under an IPTG-inducible promoter, *pilQ* was cloned in pKHT5 using a SLIC (sequence and ligation independent cloning) method(Jeong *et al*, 2012) because *pilQ* has a KpnI site in the coding sequence. Briefly, *pilQ* with a Φ10 RBS was amplified using a primer pair 5’-GCTTAGTCGACAGCTAGCC*<u>GGATCC</u>*TTTAAGAAGGAGATATACATATGAACAGTGGCCTC TCG-3’ and 5’-TTATGCTA*<u>AAGCTT</u>*GCATGCGGTACCTCAGCGACCGATTGCGATG-3’. The PCR product was mixed with pKHT105 digested with KpnI and BamHI, and processed with T4 DNA polymerase (NEB) for 2 min at RT, and transformed into *E.coli* competent cells to generate pOKP121.

#### pOKP139

To clone *pilC* under arabinose promoter, PAO1^tw^ *pilC* with native RBS was amplified using a primer pair 5’-GCTA*<u>CTGCAG</u>*CGTGACCAAGGATTAATCC-3’ and 5’-GCTA*<u>TCTAGA</u>*AGTTATCCGACGACGTTGC-3’. The PCR product was digested with EcoRI and XbaI and ligated with pJN105 digested with the same enzymes to generate pOKP139.

#### pOKP154

To construct a plasmid that expresses *slkB*-*mScarlet* under a weak IPTG-inducible promoter (*Ptoplac-dn1*), *slkB* was amplified using a primer pair 5’-GCTA*<u>CCATG**G**</u>***CT**ATCCCGCGCATTCTC-3’ and 5’-GCTA*<u>CTGCAG</u>*GCGCTCGCTTGGCTC-3’. The PCR product was digested with NcoI and PstI and ligated with pBO037 digested with the same enzymes to generate pOKP154.

### *P. aeruginosa* strain construction

#### Gene knockout and chromosomal gene fusion

Plasmids for gene knockout or gene fusion were transferred into *P. aeruginosa* by conjugation from and *E. coli* donor [Sm10(λpir)] on LB plates. Transconjugants were selected on Vogel-Bonner minimal medium (VBMM) agar supplemented with 30 μg/mL gentamicin and purified on LB agar supplemented with 30 μg/mL gentamicin. For gene deletions in the *slkAB* mutants, 10 μg/mL gentamicin was used to prevent enrichment of random mutations that suppress the barrier defect. To remove the integrated plasmid for allele exchange, purified transconjugants was grown overnight in LB with 10 μg/mL gentamicin at 37 °C. The overnight culture was diluted 1:100 in LB and grown for 4hrs at 30 °C to allow plasmid recombination to take place. The resulting culture was diluted 1:20 and 100 μL of the dilution was spread on LB agar supplemented with 5% sucrose. The plates were incubated overnight at 30 °C to select for colonies that lost the plasmid via double crossover. Sucrose-resistant colonies were patched on LB agar supplemented with either 5% sucrose or 30 μg/mL gentamicin. Gentamicin-sensitive colonies were further screened for the desired gene alteration by PCR with a primer pair that flank the altered chromosomal region.

#### Transformation of *P. aeruginosa* strains

All transformations of *P. aeruginosa* strains were performed by electroporation using electrocompetent cells made with 0.3 M sucrose solution. Briefly, 6 mL overnight culture *P. aeruginosa* strains were washed with 0.3 M sucrose solution 4 times at room temperature. After the final wash, the cell pellets were resuspended in 200 μL of 0.3 M sucrose solution. The resulting competent cells (50 μL) were electroporated with 50 ng of plasmid DNA.

#### Integration of plasmids for gene expression at the Tn7 attachment site

Plasmids for gene expression from the Tn7 attachment site were introduced into the target cell by electroporation following a previously described method (Choi & Schweizer, 2006). The recipient strain was co-electroporated with a plasmid for gene expression and pTNS3 [*bla oriR6K tnsABCD*] that expresses Tn7 transposase. Transformants were selected on LB agar supplemented with 30 μg/mL gentamicin. For gene deletions in the strains that can show an envelope permeability defect, 10 μg/mL gentamicin was used to prevent enrichment of random mutations that suppress the permeability defect. Gentamicin-resistant colonies were subject to diagnostic PCR with primer pairs 5’-CACAGCATAACTGGACTGATTTC-3’ and 5’-GCACATCGGCGACGTGCTCTC-3’ to verify the integration of the transposon at the Tn7 site.

#### OKP5 (*ΔslkA*), OKP6 (*ΔslkB*), OKP7 (*ΔslkAB*), and OKP12 (*ΔpilQ*)

To delete *slkA*, *slkB*, *slkAB*, or *pilQ* in PAO1, pOKP10, pOKP11, pOKP12, or pOKP23 was transferred to PAO1 (the twitching-defective strain) using Sm10(λpir). Candidates for the deletion strains were obtained by selection with gentamicin and counter-selection with sucrose as described above. Colonies with the desired gene deletion were screened with primer pairs 5’-TTCGGATGCTGCGGCAACTC-3’ and 5’-CAAACCAGGCGCCGAAAAGG-3’ for *ΔslkA*, 5’-GATGATCGGCACGAGAAAGG-3’ and 5’-CTTCTTCGATGATGACGTTG-3’ for *ΔslkB,* 5’-TTCGGATGCTGCGGCAACTC-3’ and 5’-CTTCTTCGATGATGACGTTG-3’ for *ΔslkAB,* 5’-ACGACTTGGCGACCTTCGTC-3’ and 5’-ATTTCGCGGTAGAGCGGGTC-3’ for *ΔpilQ*.

#### OKP13

To generate OKP13 (*ΔpilQ ΔslkAB*), *pilQ* was deleted in OKP7(*ΔslkAB*) by basically using the same procedure for gene deletion except that 10 μg/mL gentamicin was used for selection of transconjugants. As OKP7 is much more susceptible to aminoglycosides than the wild type *P. aeruginosa* strains due to its defective OM barrier function, a lower concentration of gentamicin was used to avoid selection of strains with random mutations that suppress the barrier defect.

#### OKP13(attTn7::pKHT105) and OKP13(attTn7::pOKP121)

pOKP121 that express *pilQ* from an IPTG-inducible *toplac-uv5* promoter and its empty vector plasmid were integrated into OKP13 by using the procedure described above.

#### HJP1

*slkAB* was deleted in PAO1^tw^ by following the same procedure described above after transferring pOKP11.

#### OKP14 and OKP15

*pilQ* was deleted in PAO1^tw^ and HJP1 by following the same procedure described above after transferring pOKP23 to generate OKP14 (PAO1^tw^ Δ*pilQ*) and OKP15(PAO1^tw^ Δ*slkAB* Δ*pilQ*)

#### OKP15(attTn7::pOKP121)

pOKP121 that express *pilQ* from an IPTG-inducible *toplac-uv5* promoter was integrated into OKP15 (PAO1^tw^ Δ*slkAB* Δ*pilQ*) by using the procedure described above.

#### OKP20 and OKP21

Strains that express *pilQ-mScarlet* sandwich fusion at the native *pilQ* locus in the PAO1^tw^ and HJP1 strain backgrounds were generated by using the procedure described above after transferring pOKP35 by conjugation.

#### OKP23

*pilC* was deleted in PAO1^tw^ by using the procedure described above with pOKP42.

#### OKP24

To generate OKP24 (Δ*pilC* Δ*slkAB*), *pilC* was deleted in HJP1 (Δ*slkAB*) by basically using the same procedure used to produce OKP23 except that 10 μg/mL gentamicin was used for selection of transconjugants to avoid enrichment of random mutations that suppress the barrier defect.

#### OKP23(attTn7::pKHT104) and OKP24(attTn7:: pKHT104)

To integrate *Ptoplac-dn1*-regulated empty vector into the Tn7 attachment site, pKHT104 was co-electroporated with pTNS3 into OKP23 and OKP24. A lower concentration of gentamicin (10 μg/mL) was used for selection to avoid enrichment of random mutations that suppress the barrier defect.

#### OKP26, OKP28, and OKP115

*xcpS, pscJ, or hxcS* was deleted in HJP1 by using pOKP43, pOKP44, or pOKP107 to generate double mutant strains in PAO1^tw^, Δ*slkAB* Δ*xcpS* (OKP26), Δ*slkAB* Δ*pscJ* (OKP28), Δ*slkAB* Δ*hxcS* (OKP115).

#### OKP29

*pilQ-mScarlet* sandwich fusion at the native *pilQ* locus was introduced in OKP23 (Δ*pilC*) using pOKP35 to generate OKP29 (PAO1^tw^ Δ*pilC pilQ-mScarlet^SW^*).

#### OKP44

To generate a Δ*pilC* Δ*pilQ* double mutant in the PAO1^tw^ strain background, *pilC* was deleted in OKP14 (Δ*pilQ*) by using pOKP42.

#### OKP45 and OKP96

*pilA* was deleted in HJP1 and PAO1^tw^ by using pOKP82 to generate OKP45 (PAO1^tw^ Δ*slkAB* Δ*pilA*) and OKP96 (PAO1^tw^ Δ*pilA*).

#### OKP46

*slkAB* was deleted in PA14 by using pOKP92 to generate a PA14 Δ*slkAB* strain.

#### OKP47

*slkAB* was deleted in PAK by using pOKP93 to generate a PAK Δ*slkAB* strain.

#### OKP64

*pilC* was deleted in PA14 by using pOKP94 to generate a PA14 Δ*pilC* strain.

#### OKP48

*pilC* was deleted in PAK by using pOKP95 to generate a PAK Δ*pilC* strain.

#### OKP49 and OKP50

*pilMNOPQ* was deleted in PAO1^tw^ and HJP1 using pOKP88 to generate OKP49 (PAO1^tw^ Δ*pilMNOPQ*) and OKP50 (PAO1^tw^ Δ*slkAB* Δ*pilMNOPQ*), respectively.

#### PAO1^tw^(attTn7::pKHT105), OKP49(attTn7::pKHT105), OKP49(attTn7::pOKP121), OKP50(attTn7::pKHT105), OKP50(attTn7::pOKP121)

pOKP121 (*Ptoplac-uv5*-regulated *pilQ*) or its empty vector plasmid pKHT105 were integrated into the Tn7 attachment site by co-electroporation with pTNS3. Candidates with the integrated plasmids were obtained by selection with 10 μg/mL gentamicin to avoid enrichment of random mutations that suppress the barrier defect.

#### OKP51

*pilQ* was deleted in PAO1^tw^ Δ*slkAB* Δ*pilC* strain by using pOKP23 to generate OKP24 (PAO1^tw^ Δ*slkAB* Δ*pilC* Δ*pilQ*). A lower concentration of gentamicin (10 μg/mL) was used for selection of transconjugants to avoid enrichment of random mutations that suppress the barrier defect.

#### HJP1(attTn7::pOKP154), OKP15(attTn7::pOKP154), OKP24(attTn7::pOKP154), OKP51(attTn7::pOKP154)

To integrate a plasmid expressing *slkB-mScarlet* from *Ptoplac-dn1* (pOKP154) into the Tn7 attachment site, pOKP154 was co-electroporated with pTNS3 into HJP1, OKP15, OKP24, OKP51. Candidates for the integrated strains were obtained by selection with 10 μg/mL gentamicin to avoid enrichment of random mutations that suppress the barrier defect.

#### OKP52 and OKP53

*fimV* or *tsaP* was deleted in OKP24 (PAO1^tw^ Δ*slkAB* Δ*pilC*) by using pOKP98 (*fimV* deletion) or pOKP99 (*tsaP* deletion) to generate OKP52 and OKP53, respectively. A lower concentration of gentamicin (10 μg/mL) was used for selection of transconjugants to avoid enrichment of random mutations that suppress the barrier defect.

#### OKP57

*pilC* was deleted in OKP47 (PAK Δ*slkAB*) by using pOKP95.

#### OKP59, OKP60, OKP61, and OKP120

*slkA-mScarlet* with a *slkB* deletion was introduced at the *slkAB* locus of PAO1^tw^, OKP14 (PAO1^tw^ Δ*pilQ*), OKP23 (PAO1^tw^ Δ*pilC*), and OKP44 (PAO1^tw^ Δ*pilC* Δ*pilQ*) strains by using pOKP101.

#### OKP62

*oprM* was deleted in PAO1 by using pOKP103.

#### OKP65

*pilC* was deleted in OKP46 (PA14 Δ*slkAB*) by using pOKP94. Candidates for the deletion strain were obtained by selection with 10 μg/mL gentamicin to avoid enrichment of random mutations that suppress the barrier defect.

#### OKP68

*pilMNOPQ* was deleted in HJP1 (PAO1 Δ*slkAB*) using pOKP88.

#### OKP69

*pilQ* was deleted in OKP45 (PAO1^tw^ Δ*slkAB* Δ*pilA*) by using pOKP23. Candidates for the deletion strain were obtained by selection with 10 μg/mL gentamicin to avoid enrichment of random mutations that suppress the barrier defect.

#### OKP68(attTn7::pKHT105) and OKP68(attTn7::pOKP121)

A plasmid expressing *pilQ* from *Ptoplac-uv5* (pOKP121) and its empty vector plasmid (pKHT105) were integrated at the Tn7 attachment site of OKP68 by co-electroporation with pTNS3. Candidates for the integrated strains were obtained by selection with 10 μg/mL gentamicin to avoid enrichment of random mutations that suppress the barrier defect.

#### OKP73, OKP74, and OKP86

*tsaP was deleted* was in PAO1^tw^, HJP1 (PAO1^tw^ Δ*slkAB*), or OKP23 (PAO1^tw^ Δ*pilC*) by using pOKP99. Candidates for the deletion strains were obtained by selection with 10 μg/mL gentamicin to avoid enrichment of random mutations that suppress the barrier defect.

#### OKP80

*pilQ* was deleted in OKP65 (PA14 *ΔslkAB ΔpilC*) by using pOKP116. Candidates for the deletion strains were obtained by selection with 10 μg/mL gentamicin to avoid enrichment of random mutations that suppress the barrier defect.

#### OKP82

*pilQ* was deleted in OKP57 (PAK *ΔslkAB ΔpilC*) by using pOKP57. Candidates for the deletion strains were obtained by selection with 10 μg/mL gentamicin to avoid enrichment of random mutations that suppress the barrier defect.

#### OKP87

*pilQ* was deleted in OKP73 (PAO1^tw^ Δ*slkAB* Δ*tsaP*) by using pOKP23. Candidates for the deletion strain were obtained by selection with 10 μg/mL gentamicin to avoid enrichment of random mutations that suppress the barrier defect.

#### OKP77, OKP78, and OKP97

*pilY1* was deleted in PAO1^tw^, HJP1 (PAO1^tw^ Δ*slkAB*), and OKP15 (PAO1^tw^ Δ*slkAB* Δ*pilQ*) by using pOKP115. Candidates for the deletion strains were obtained by selection with 10 μg/mL gentamicin to avoid enrichment of random mutations that suppress the barrier defect.

### Generation of the transposon insertion library

*P. aeruginosa* strains were mutagenized with a mariner transposon delivery vector, pBTK30, (Goodman *et al*, 2004) using the following mating protocol. The donor strain SM10(*λpir*) carrying pBTK30 was grown in LB supplemented with 50 μg/mL ampicillin at 37 °C and *P. aeruginosa* strains were grown in LB at 42 °C overnight. The next morning, cultures were concentrated and adjusted to an OD600 of 5 for the donor and 10 for the recipient. Equal volumes of donor and recipient were mixed together and 50 μL aliquots were spotted on prewarmed LB plates. Mating was allowed to proceed at 37 °C for one hour prior to resuspension in VBMM supplemented with 30 μg/mL gentamicin. Transconjugants were selected on VBMM agar supplemented with 30 μg/mL gentamicin at 30 °C for 24 hours. Donor-only and recipient-only controls were performed in parallel to ensure proper selection of *P. aeruginosa* transconjugants. More than 5 million transconjugant colonies were collected from agar plates by suspension in VBMM broth supplemented with 30 μg/mL gentamicin and frozen at -80 °C.

### Transposon sequencing

A transposon mutant library of PAO1 was thawed and incubated in LB for roughly two doublings. The resuscitated mutant library culture was diluted in LB either lacking or containing 10 μg/mL erythromycin and incubated for 10 doublings to a final OD600 of 0.5 at 37 °C. After the incubation, the cells were pelleted and frozen. Genomic DNA from each cell pellet was extracted, fragmented, and poly-C tailed as previously described (Lai *et al*, 2017). The transposon-chromosome junctions in the resulting DNA were amplified by using Easy-A Hi-Fi Cloning System (Agilent Technologies). The primers used were the poly-C tail-specific primer 5’-GTGACTGGAGTTCAGACGTGTGCTCTTCCGATCTGGGGGGGGGGGGGGGG-3’ and the transposon-specific primer 5’-GGTTCTGGACCAGTTGCGTGAG-3’. The transposon-chromosome junctions were further amplified in a second, nested PCR with the primers that add sequencing barcodes to each mutant library, NEBNext Multiplex Oligos for Illumina (NEB), and the transposon-specific primer 5’-AATGATACGGCGACCACCGAGATCTACACTCTTTGTTTTCTGGAAGGCGAGCATCGTTTG-3’. The final PCR products were quantified and equal amounts of each barcoded library were mixed. The pooled sequencing library was run on a 2% agarose gel, and DNA fragments ranging from 200 and 500 bp were excised and purified using QIAquick Gel Extraction Kit (Qiagen). The resulting library was sequenced using a MiSeq reagent kit V3 (150-cycle) (Illumina) with the custom primer 5’-CTAGAGACCGGGGACTTATCAGCCAACCTGTTA-3’. Sequencing reads were trimmed using trimmomatic (Bolger *et al*, 2014) to remove adaptor sequences, and mapped to chromosomal TA dinucleotides (mariner insertion sites) on the *P. aeruginosa* PAO1 genome (NC_002516) using bowtie 1.0.0 (Langmead *et al*, 2009). Differences in the total number of reads at any given TA site between untreated and erythromycin-treated samples were determined using a Mann-Whitney U test. Transposon insertion profiles were visualized using the Sanger Artemis Genome Browser and Annotation tool.

### Whole genome sequencing

The genomic DNA of PAO1 and PAO1^tw^ strains was extracted from overnight grown cultures using Wizard^®^ Genomic DNA Purification Kit (Promega) according to the manufacturer’s manual. Purification of the extracted genomic DNA was performed using Genomic DNA Clean & Concentrator™-25 (Zymo Research) according to the manufacturer’s manual. The genomic DNA samples were sequenced by Macrogen Inc. using an Illumina HiSeq platform. Sequencing reads were analyzed with Geneious Prime software using PAO1 genome (NC_002516) as a reference.

### Suppressor selection

To select for suppressors of the erythromycin susceptibility phenotype of OKP7 (PAO1 *ΔslkAB*), the strain was mutagenized by random insertion of a mariner-based transposon from pBTK30, as described (Goodman *et al*, 2004). The mutant library was spread on LB agar supplemented with 50 μg/mL erythromycin to select for mutants that can suppress the OM permeability defect. The transposon insertion sites of the suppressors were determined by arbitrarily primed PCR. The first round was performed with a primer pair 5’-GGCCACGCGTCGACTAGTACNNNNNNNNNNGATAT-3’ and 5’-GGTTCTGGACCAGTTGCGTGAG-3’. The second, nested PCR was performed with 5’-GGCCACGCGTCGACTAGTAC-3’ and 5’-CGAACCGAACAGGCTTATGTCAATTCG-3’ to increase specificity and sensitivity. The resulting PCR products were sequenced with the primer 5’-CGAACCGAACAGGCTTATGTCAATTCG-3’ that anneal to the transposon sequence to identify the transposon-chromosome junctions.

### Twitching motility assay

To examine type IV pili-mediated twitching motility, a single colony from a freshly streaked LB agar plate was picked with a toothpick and stab-inoculated through LB agar (1% agar) to the polystyrene dish. After incubation for 48 hours at 30 °C, the LB agar was removed and the cells attached to the polystyrene dish were stained with 1% crystal violet for 5 min. The dish was then rinsed with water to remove excess stain and the stained zone was photographed after air drying.

### Phylogenetic tree generation

To generate the phylogenetic tree, the amino acids sequence of PA5122 and PA5123 were input into BLASTp and searched against the NCBI “non redundant” (*nr*) database with an *e*-value cutoff of 1e^-6^ for each protein (Pruitt *et al*, 2005). We used a complex and diverse set of 1773 bacterial taxa called ‘Representative Genomes’ that is available on NCBI (ftp://ftp.ncbi.nlm.nih.gov/blast/db/, Representative_Genomes.00.tar.gz). The phylogenetic tree was constructed using phyloT (http://phylot.biobyte.de/) and BLASTp results were plotted against the tree. The tree was visualized and annotated using iTOL (http://itol.embl.de/) (Letunic & Bork, 2016).

---

**Figure S1.**
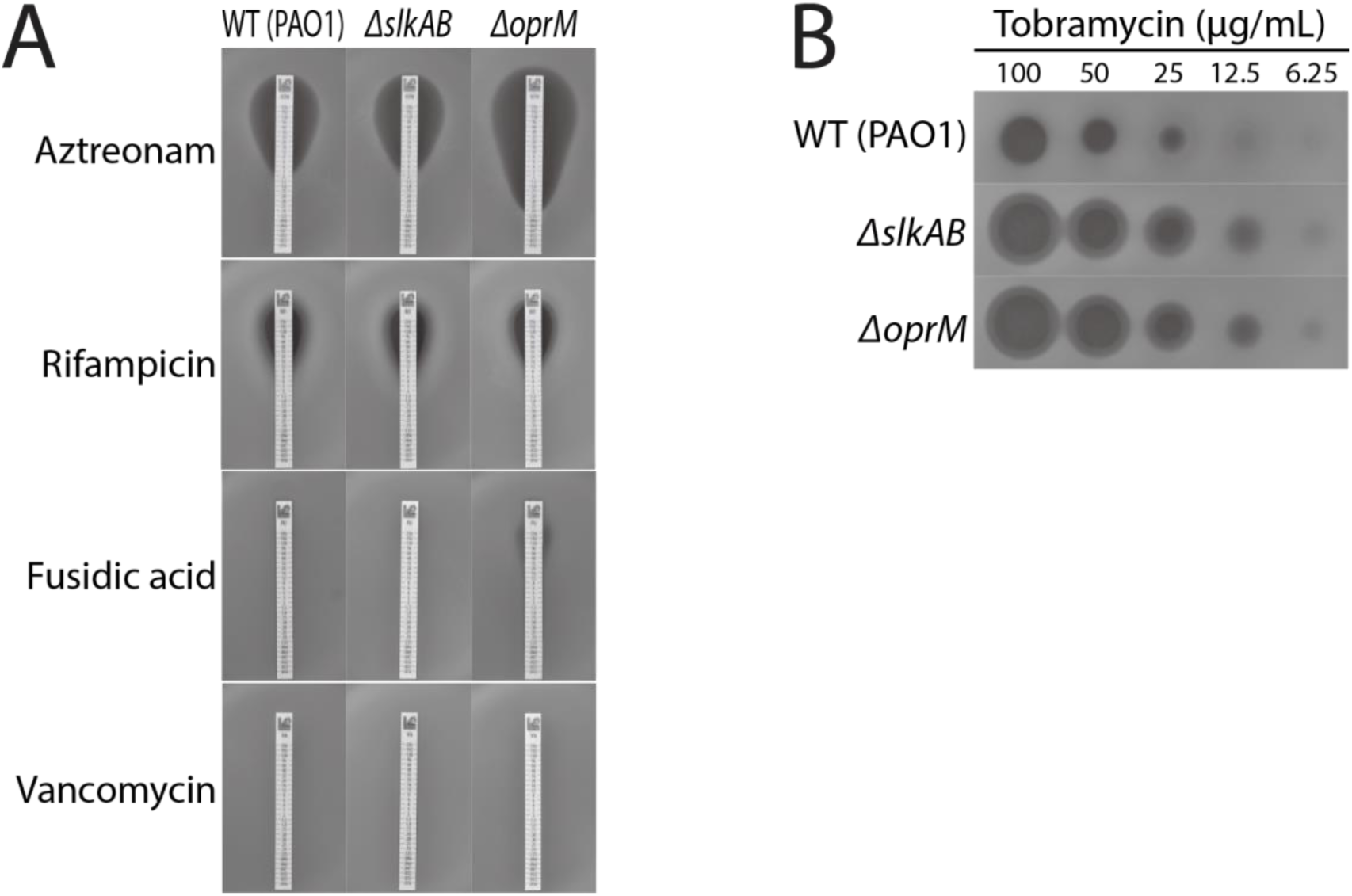
Comparison of the sensitivity of the PAO1, OKP7 (*ΔslkAB*), and OKP62 (*ΔoprM*) strains to various antibiotics. **(A)** Lawns of the PAO1 and its indicated mutant derivatives were plated in soft agar, incubated with indicated antibiotic test strips for 24 hours at 37 °C, and imaged as in Figure 1E**. (B)** Lawns of the same *P. aeruginosa* strains were spotted with 5 μL aliquots of tobramycin solutions at the indicated concentrations and imaged after incubation for 24 hours at 37 °C.

**Figure S2.**
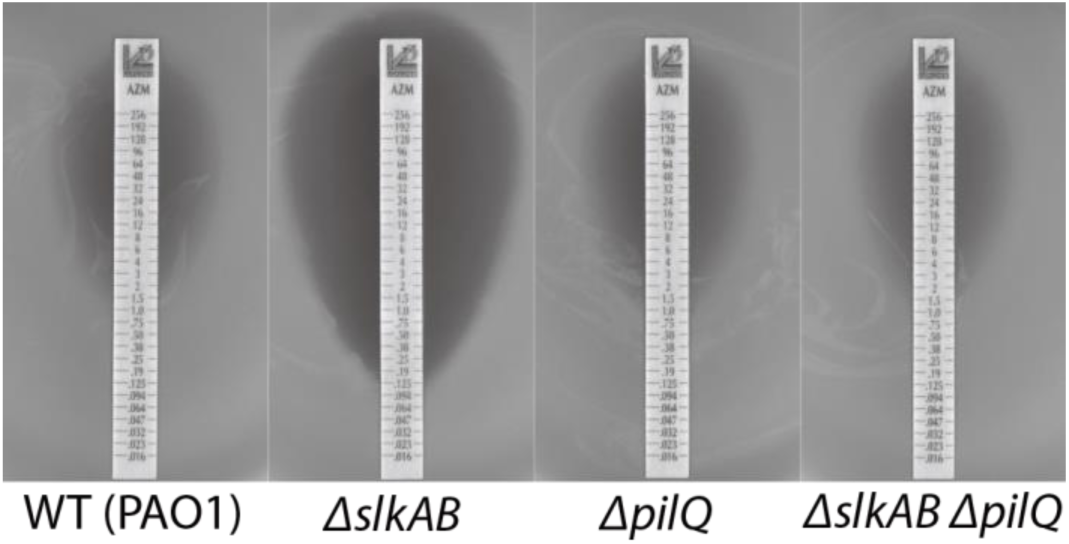
Azithromycin sensitivity of the *ΔslkAB* strain is suppressed by *ΔpilQ*. Lawns of PAO1 and its indicated mutant derivatives, OKP7(*ΔslkAB*), OKP14(*ΔpilQ*), and OKP15 (*ΔslkAB ΔpilQ*), were plated in soft agar, incubated with azithromycin test strips for 24 hours at 37 °C, and imaged as in Figure 2C.

**Figure S3.**
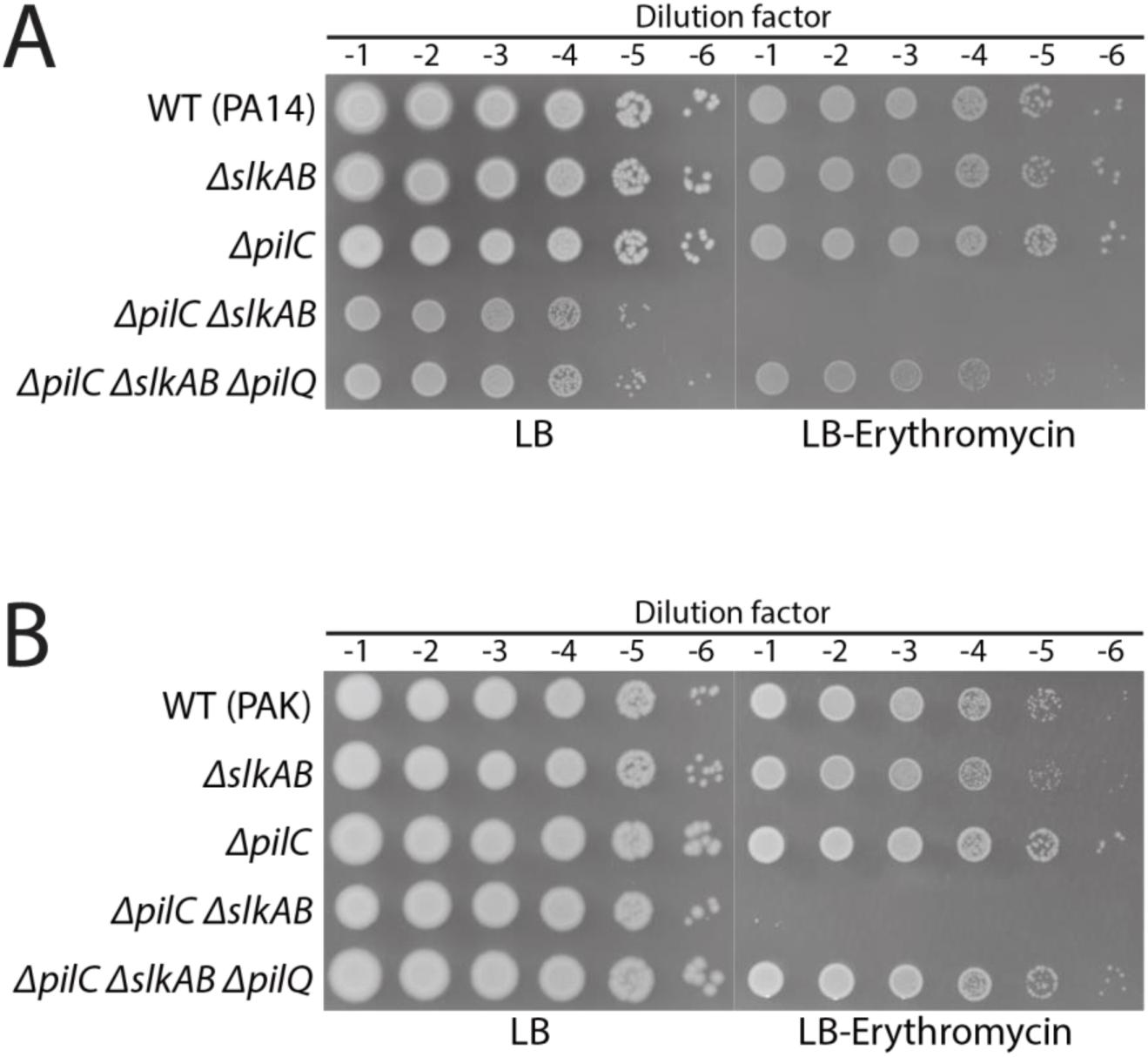
Inactivation of the IM complex assembly causes an OM secretin channel-dependent permeability barrier defect in the *ΔslkAB* mutant of PA14 (A) and PAK (B) strains. **(A)** PA14 and its mutant derivatives, OKP46 (*ΔslkAB*), OKP64 (Δ*pilC*), OKP65 (Δ*pilC ΔslkAB*), and OKP80 (Δ*pilC ΔslkAB* Δ*pilQ*), were grown overnight in LB, serially diluted, spotted onto LB agar lacking or containing 50 μg/mL erythromycin, and imaged after incubation for 20 hours at 37 °C. **(B)** Erythromycin susceptibility of PAK, and its mutant derivatives, OKP47 (*ΔslkAB*), OKP48 (Δ*pilC*), OKP57 (Δ*pilC ΔslkAB*), and OKP82 (Δ*pilC ΔslkAB* Δ*pilQ*), was examined in the same way.

**Figure S4.**
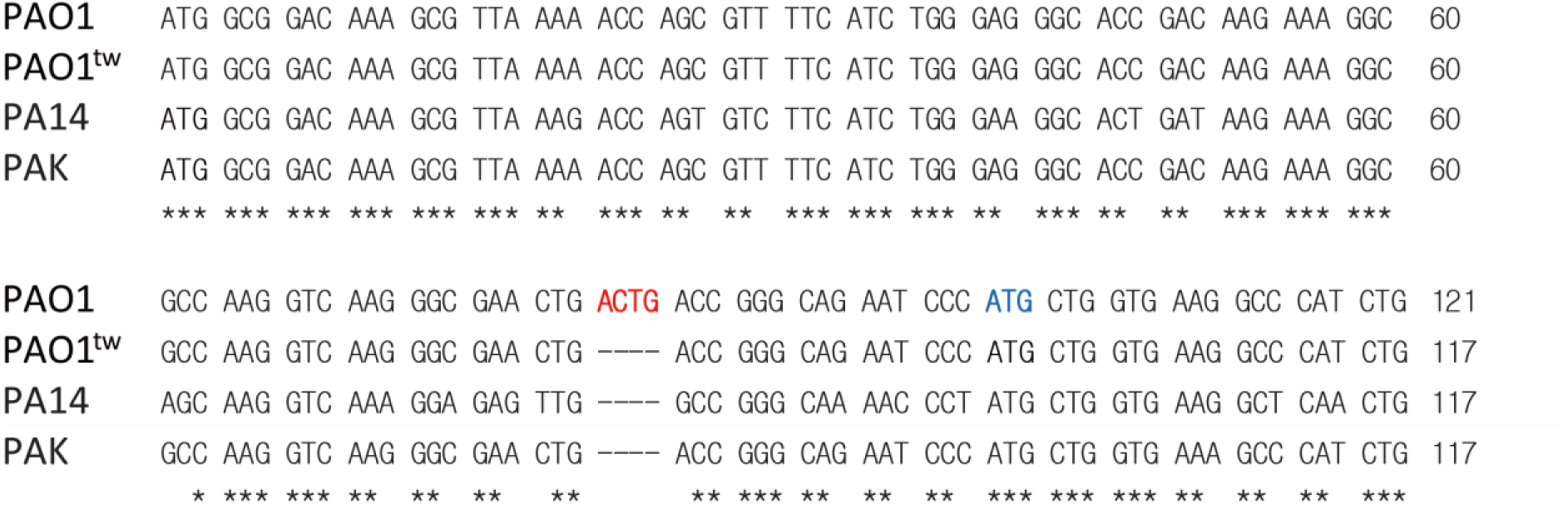
The PAO1 strain used for the Tn-Seq and initial characterization has a frameshift mutation by 4 base insertion in the coding sequence of *pilC*. Alignment of the *pilC* coding sequences of several *P. aeruginosa* strains revealed that the PAO1 strain we initially used has a frameshift mutation in the *pilC* gene by insertion of 4 bases (indicated in red) after the 27^th^ codon. The same frameshift mutation was present in the PAO1 reference genome (NC_002516), but was not recognized as a frameshift mutation because an internal ATG codon (indicated in blue) was misannotated as the start codon. Twitching proficient strains, PAO1^tw^, PA14, and PAK, do not have this insertion. Conserved bases among the *pilC* coding sequences of different *P. aeruginosa* strains are indicated as * at the bottom.

**Figure S5.**
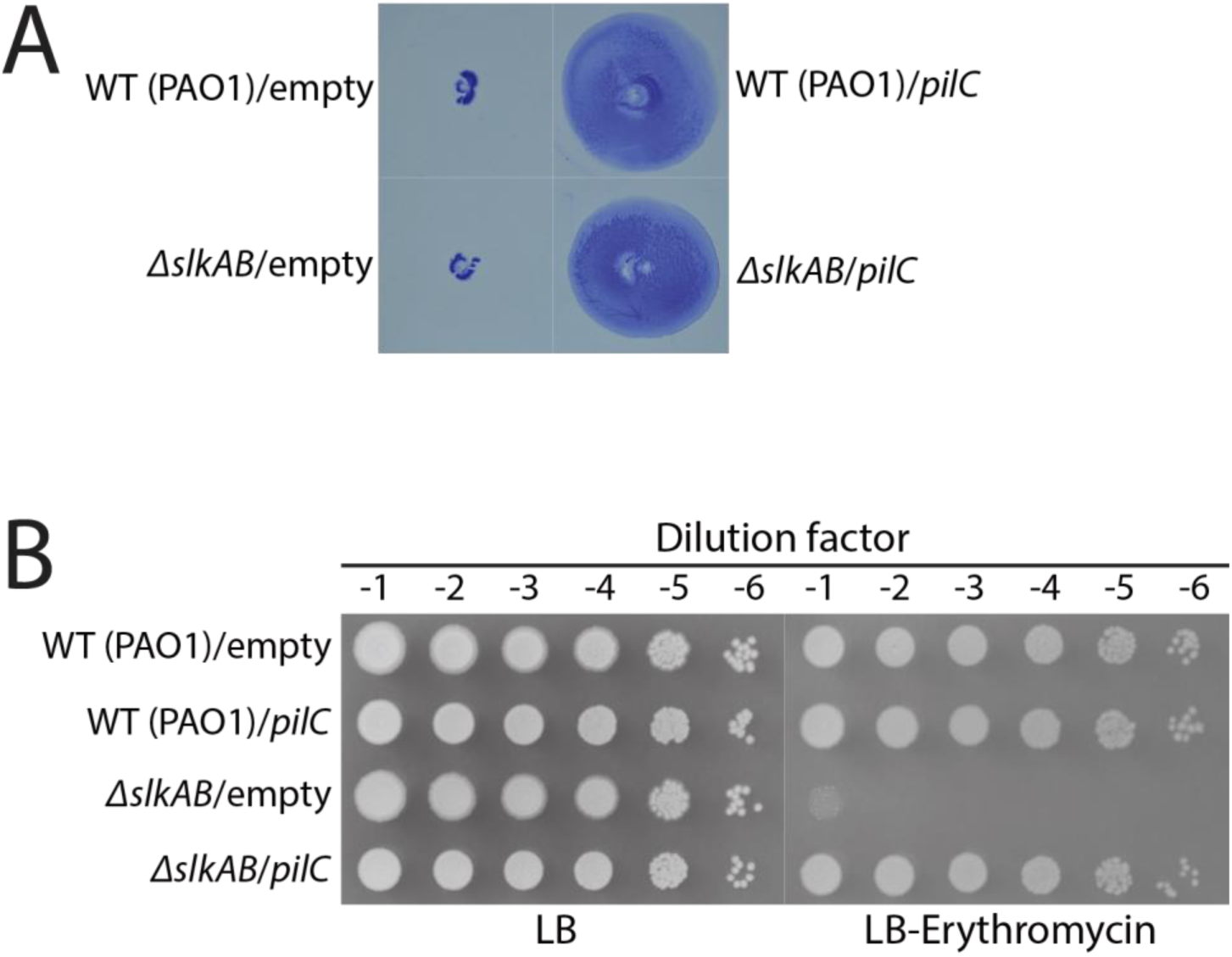
Expression of the WT *pilC* gene not only restores the twitching motility of PAO1 and its *ΔslkAB* derivative, but also suppresses erythromycin sensitivity of the *ΔslkAB* derivative. **(A)** The twitching-defective PAO1 strain and its *ΔslkAB* derivative (OKP7) harboring pJN105 (empty vector) or pOKP139 expressing *pilC* from the twitching-proficient strain were tested for twitching motility on the polystyrene dish as described in Fig. 3A. **(B)** The same strains were grown overnight in LB supplemented with 10 μg/mL gentamicin, serially diluted in LB, spotted onto LB agar lacking or containing 50 μg/mL erythromycin, and imaged after incubation for 20 hours at 37 °C.

**Figure S6.**
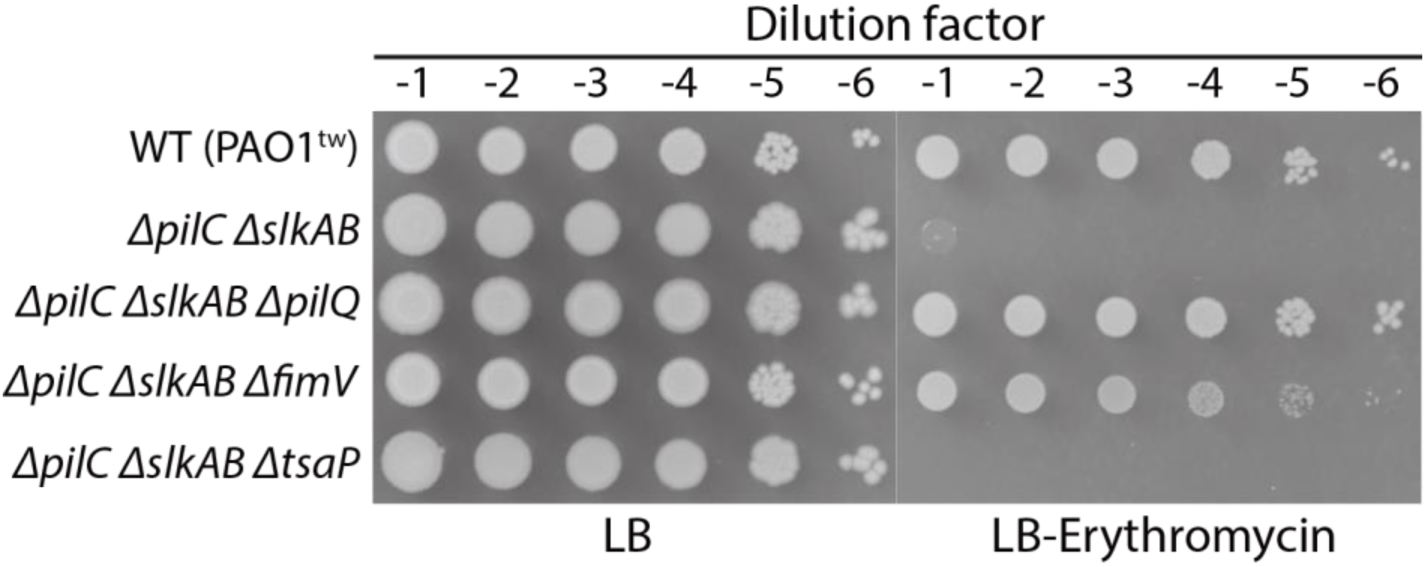
Inactivation of TsaP does not suppress the synthetic envelope defect of the *ΔpilC ΔslkAB* strain. PAO1^tw^ (WT) and its mutant derivatives, OKP24 (*ΔpilC ΔslkAB*), OKP51 (*ΔpilC ΔslkAB ΔpilQ*), OKP52 (*ΔpilC ΔslkAB ΔfimV*), and OKP53 (*ΔpilC ΔslkAB ΔtsaP*), were grown overnight, serially diluted, spotted onto LB agar lacking or containing 50 μg/mL erythromycin, and imaged after incubation for 20 hours at 37 °C.

**Figure S7.**
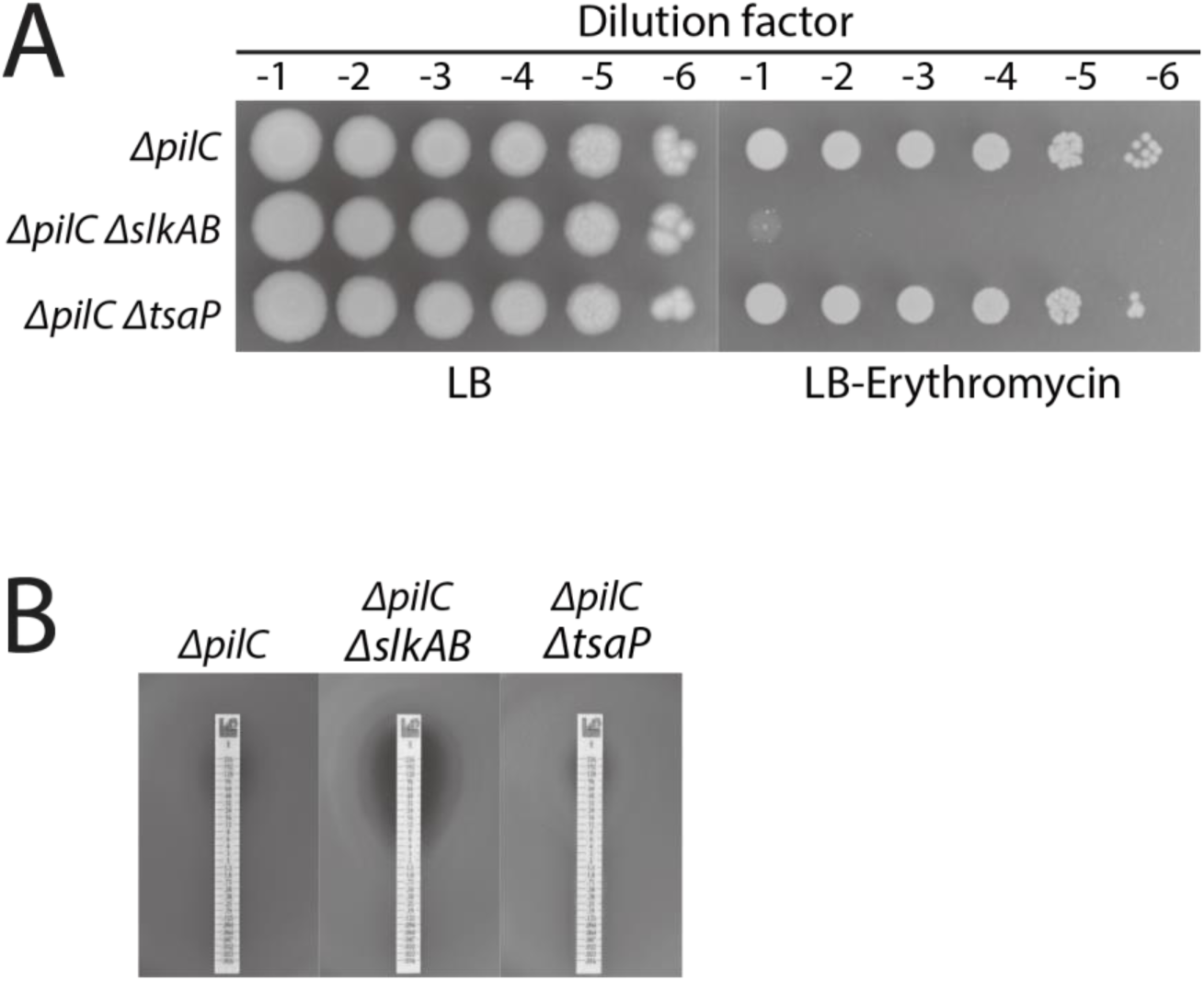
Inactivation of TsaP does not cause a synthetic envelope defect in the *ΔpilC* strain. **(A)** Mutant derivatives of PAO1^tw^, OKP23 (*ΔpilC*), OKP24 (*ΔpilC ΔslkAB*), and OKP74 (*ΔpilC ΔtsaP*), were grown overnight, serially diluted, spotted onto LB agar lacking or containing 50 μg/mL erythromycin, and imaged after incubation for 20 hours at 37 °C. **(B)** Lawns of the same *P. aeruginosa* strains were plated in soft agar, incubated with erythromycin test strips for 24 hours at 37 °C, and photographed.

**Figure S8.**
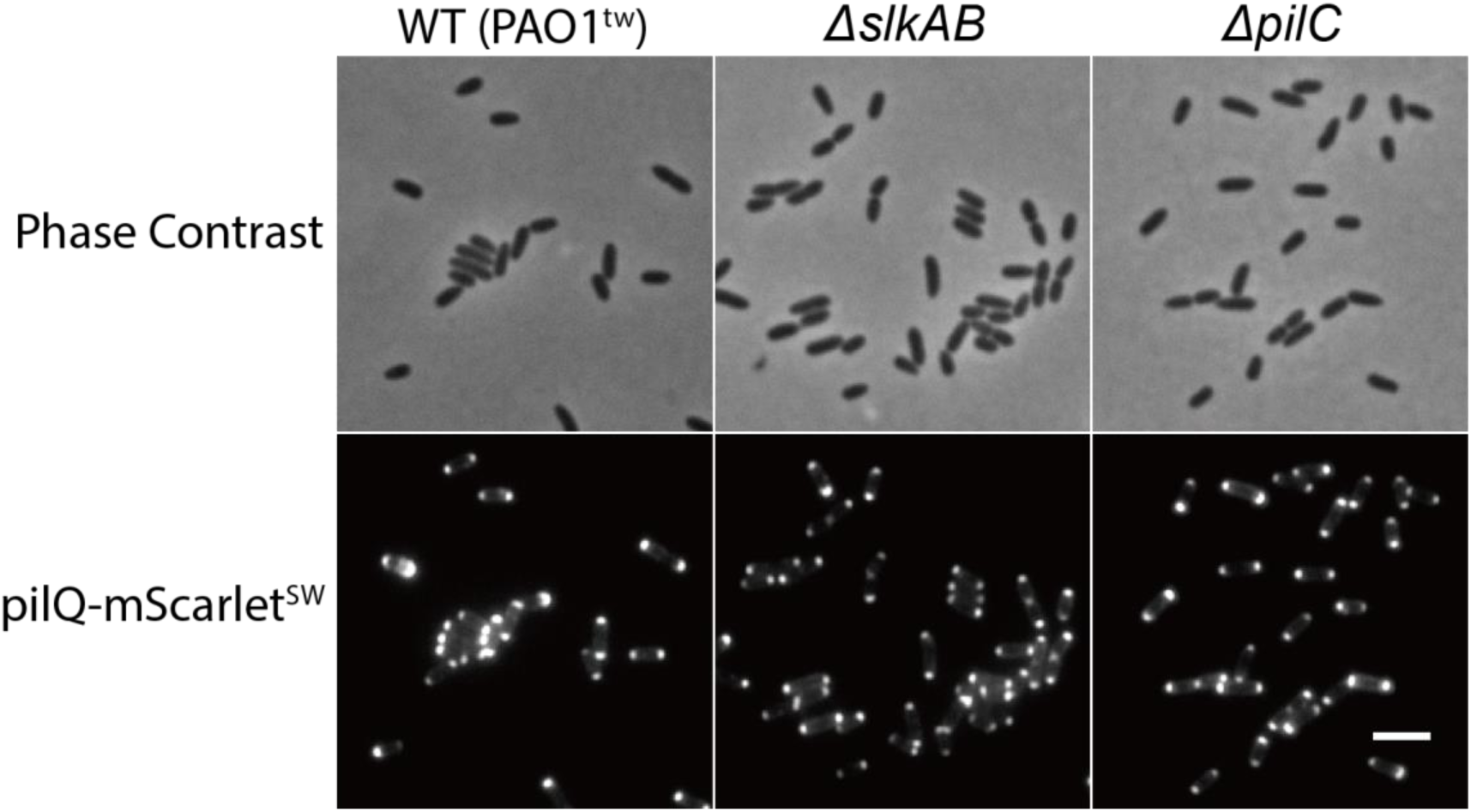
PilQ localizes to the poles. PAO1^tw^ and its *ΔslkAB* and *ΔpilC* mutant derivatives that express *pilQ-mScarlet^SW^* from its native locus (OKP20, OKP21, and OKP29) were grown overnight in LB. The overnight cultures were diluted 1:100 in M9-glucose (0.2%) medium, grown at 37 °C to an exponential phase (OD600 = 0.2∼0.3), and imaged on 2% agarose pads containing 1X M9 salts. Bar equals 3 μm.

**Figure S9.**
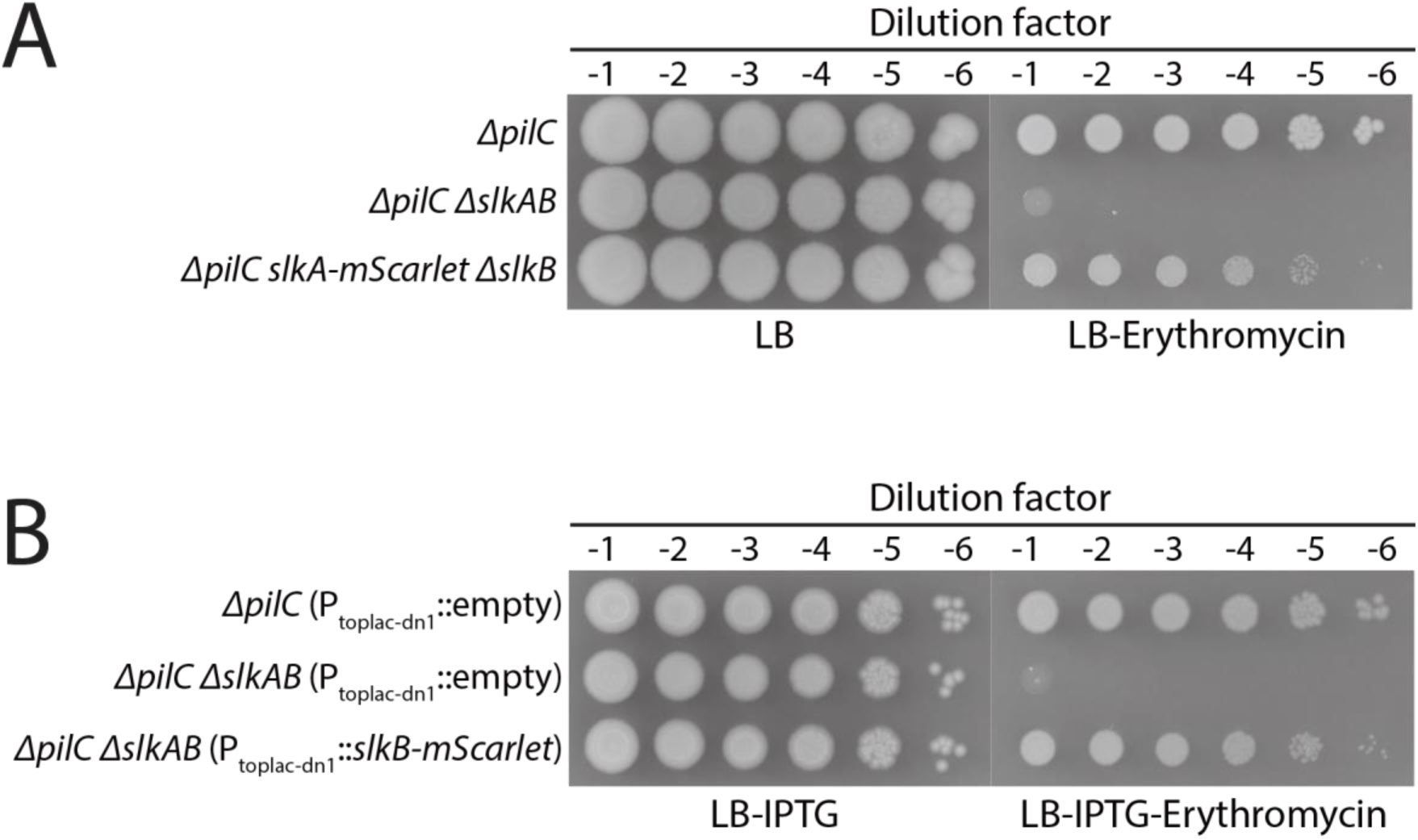
SlkA-mScarlet and SlkB-mScarlet are partially functional. **(A)** OKP23 [PAO1^tw^ Δ*pilC*], OKP24 [PAO1^tw^ Δ*slkAB* Δ*pilC*], and OKP61 [PAO1^tw^ Δ*pilC slkA-mScarlet* Δ*slkB*] strains were grown overnight in LB, serially diluted, and spotted on LB agar lacking or containing 50 μg/mL erythromycin. The plates were photographed after incubation for 28 hours at 37 °C to visualize suppression of the barrier defect by *slkA-mScarlet* expression. **(B)** OKP23 (attTn7::pKHT104, empty vector), OKP24 (attTn7::pKHT104), and OKP24 (attTn7::pOKP154, Ptoplac-dn1::*slkB-mScarlet*) strains were grown overnight in LB, serially diluted, and spotted on LB agar supplemented with 100 μM IPTG or 100 μM IPTG and 50 μg/mL erythromycin. The plates were photographed after incubation for 20 hours at 37 °C.

**Figure S10.**
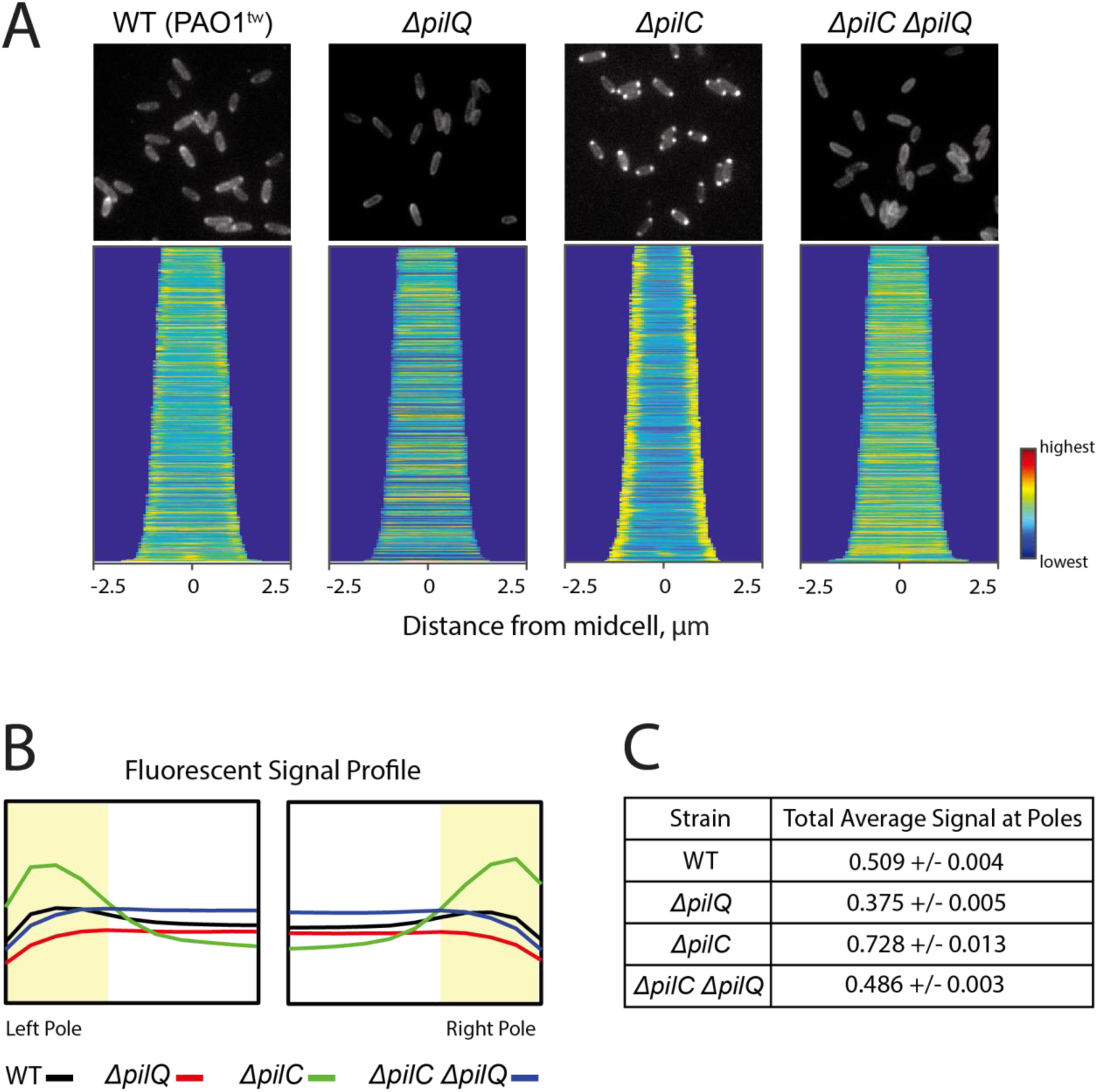
PilQ-dependent polar localization of SlkB and its enhancement upon inactivation of T4P platform PilC. PAO1^tw^ derivatives that express *slkB*-mScarlet under an IPTG-inducible toplac-dn1 promoter, HJP1[PAO1^tw^ Δ*slkAB*](attTn7::pKOP154), OKP15[PAO1^tw^ Δ*slkAB* Δ*pilQ*](attTn7::pKOP154), OKP24[PAO1^tw^ Δ*slkAB* Δ*pilC*](attTn7::pKOP154), and OKP51[PAO1^tw^ Δ*slkAB* Δ*pilC* Δ*pilQ*](attTn7::pKOP154) were grown overnight in LB. The overnight cultures were diluted 1:100 in M9-glucose (0.2%) medium supplemented with 100 μM IPTG, grown to an exponential phase (OD600 = 0.2∼0.3), and imaged on 2% agarose pads containing 1X M9 salts. The images were processed and presented in the same way as described for Figure 4.

**Figure S11.**
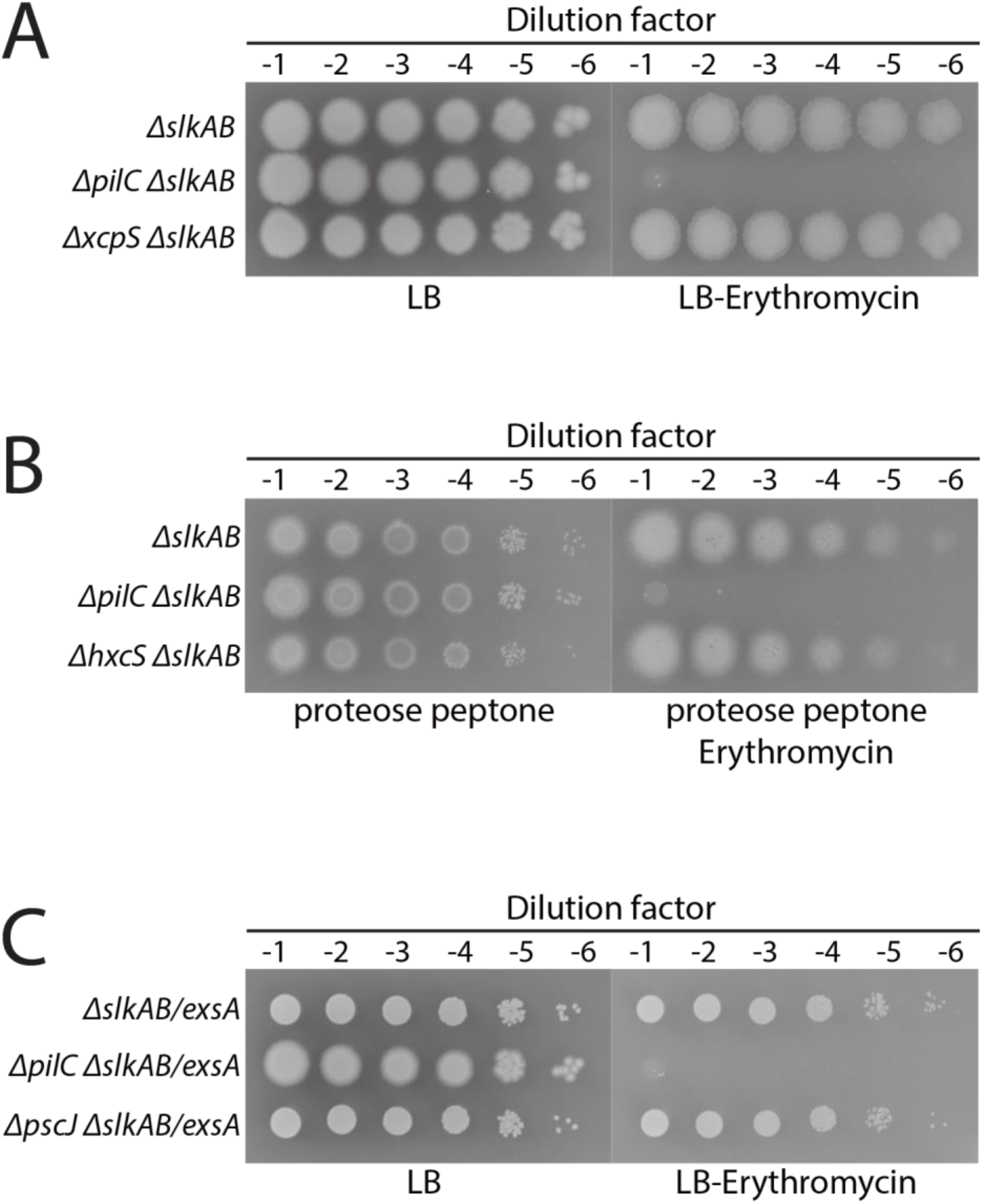
Slk proteins are likely to function specifically for T4PS in *P. aeruginosa*. **(A)** Mutant derivatives of PAO1^tw^, HJP1 (PAO1^tw^ Δ*slkAB*), OKP24 (PAO1^tw^ Δ*slkAB* Δ*pilC*), and OKP26 (PAO1^tw^ Δ*slkAB* Δ*xcpS*) were grown overnight in LB, serially diluted, and spotted on LB agar lacking or containing 50 μg/mL erythromycin. The plates were photographed after incubation for 20 hours at 37 °C. **(B)** HJP1, OKP24, and OKP115 (PAO1^tw^ Δ*slkAB* Δ*hxcS*) were grown overnight in LB, diluted on phosphate-limiting proteose peptone medium, and spotted on proteose peptone agar lacking or containing 50 μg/mL erythromycin for induction of Hxc T2SS genes. **(C)** HJP1, OKP24, and OKP28 (PAO1^tw^Δ*slkAB* Δ*pscJ*) harboring pBO48 that express *exsA* for induction of T3SS genes were grown overnight in LB supplemented with 10 μg/mL gentamicin, serially diluted in LB, and spotted on LB agar lacking or containing 50 μg/mL erythromycin.

**Figure S12.**
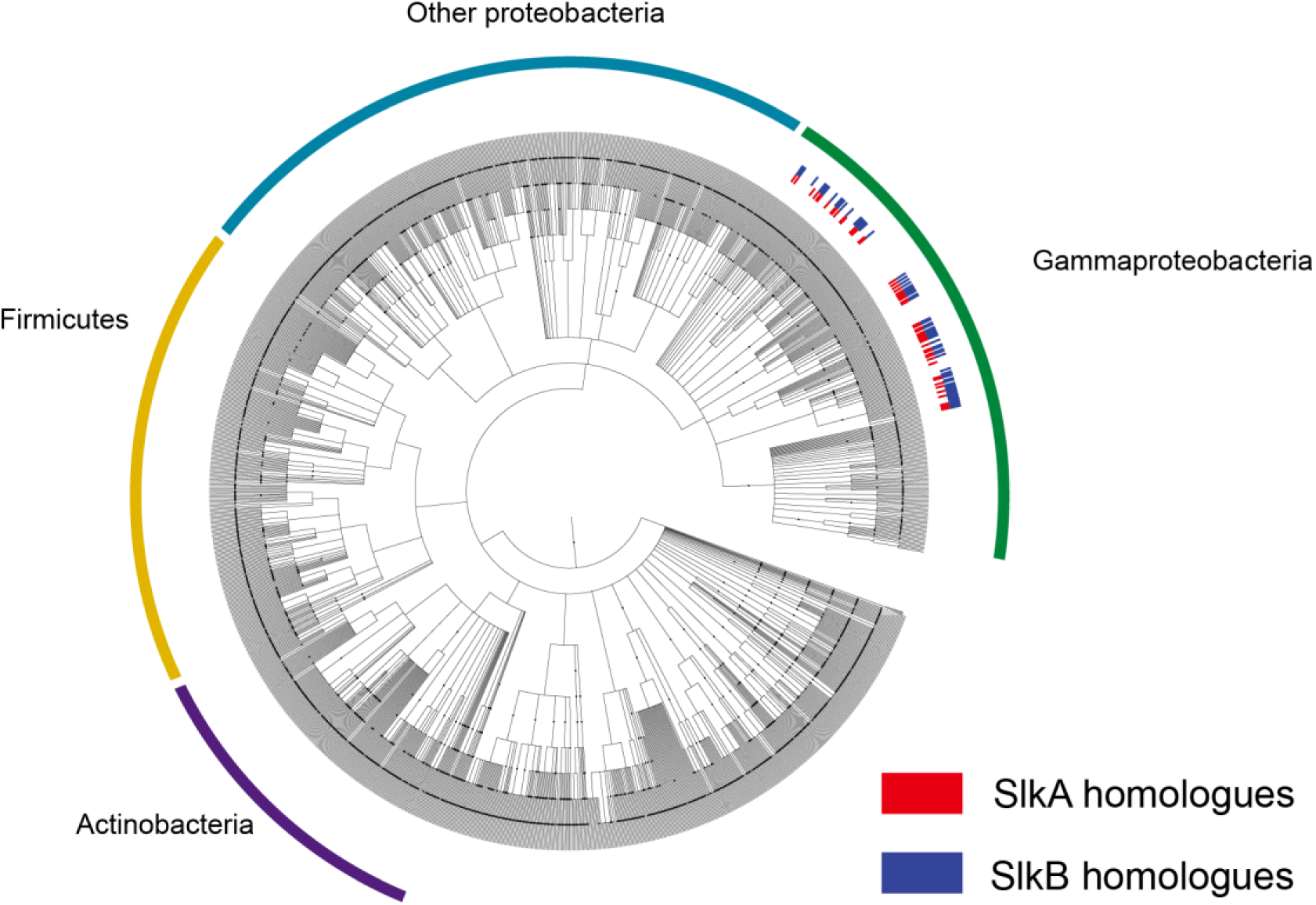
Phylogenetic distribution of SlkA and SlkB homologues. Shown is a phylogenetic tree depicting the occurrence of SlkA (red) and SlkB (blue) homologues among bacteria. The phylogenetic tree was constructed using phyloT (http://phylot.biobyte.de/) and visualized and annotated using iTOL (http://itol.embl.de/).

**Figure S13.**
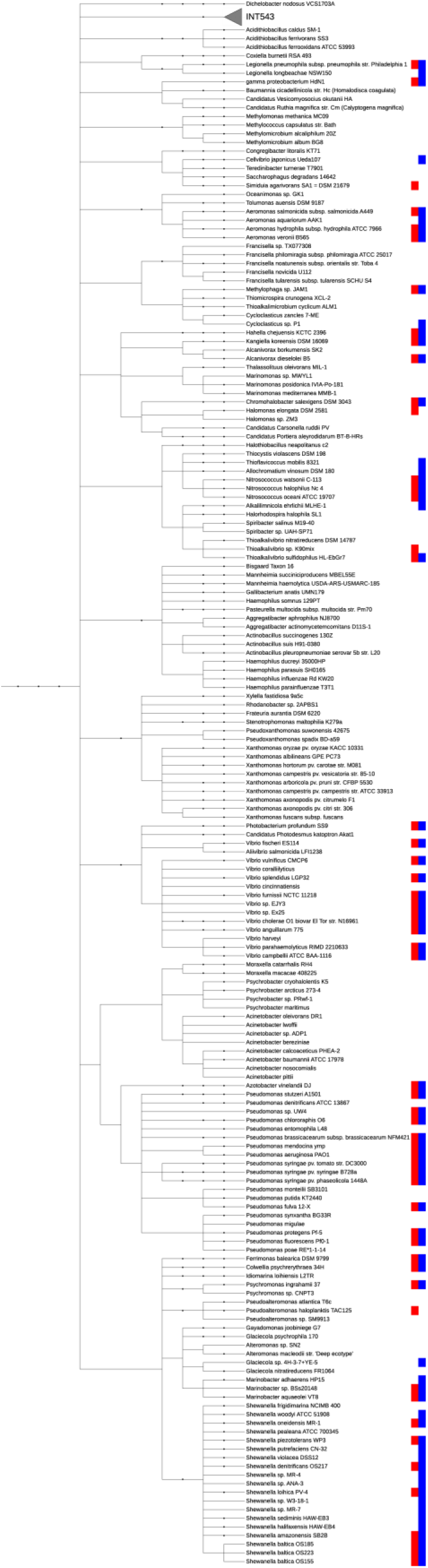
Phylogenetic tree showing the distribution of SlkA and SlkB homologues among gamma-proteobacteria

**Table S3.**
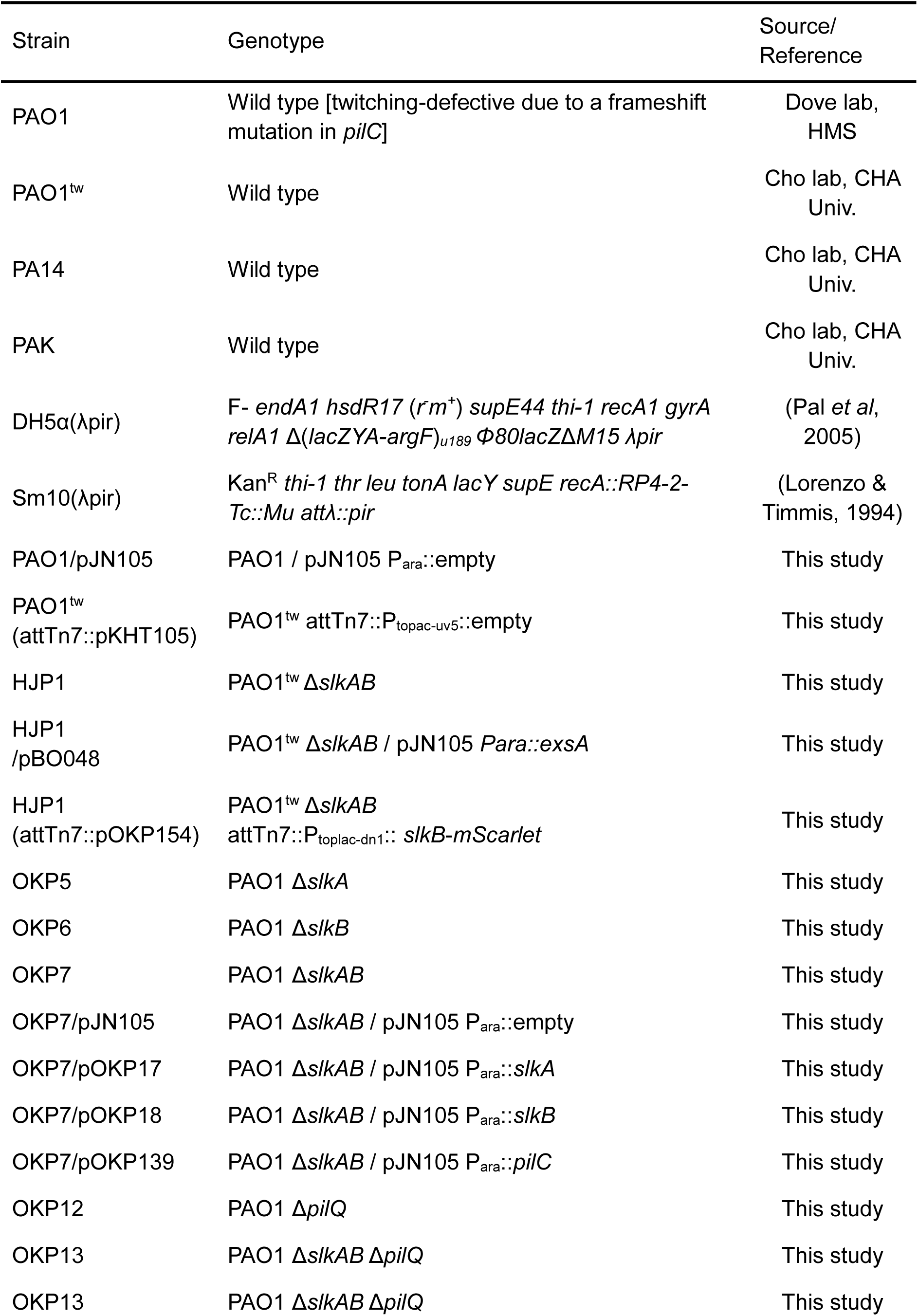
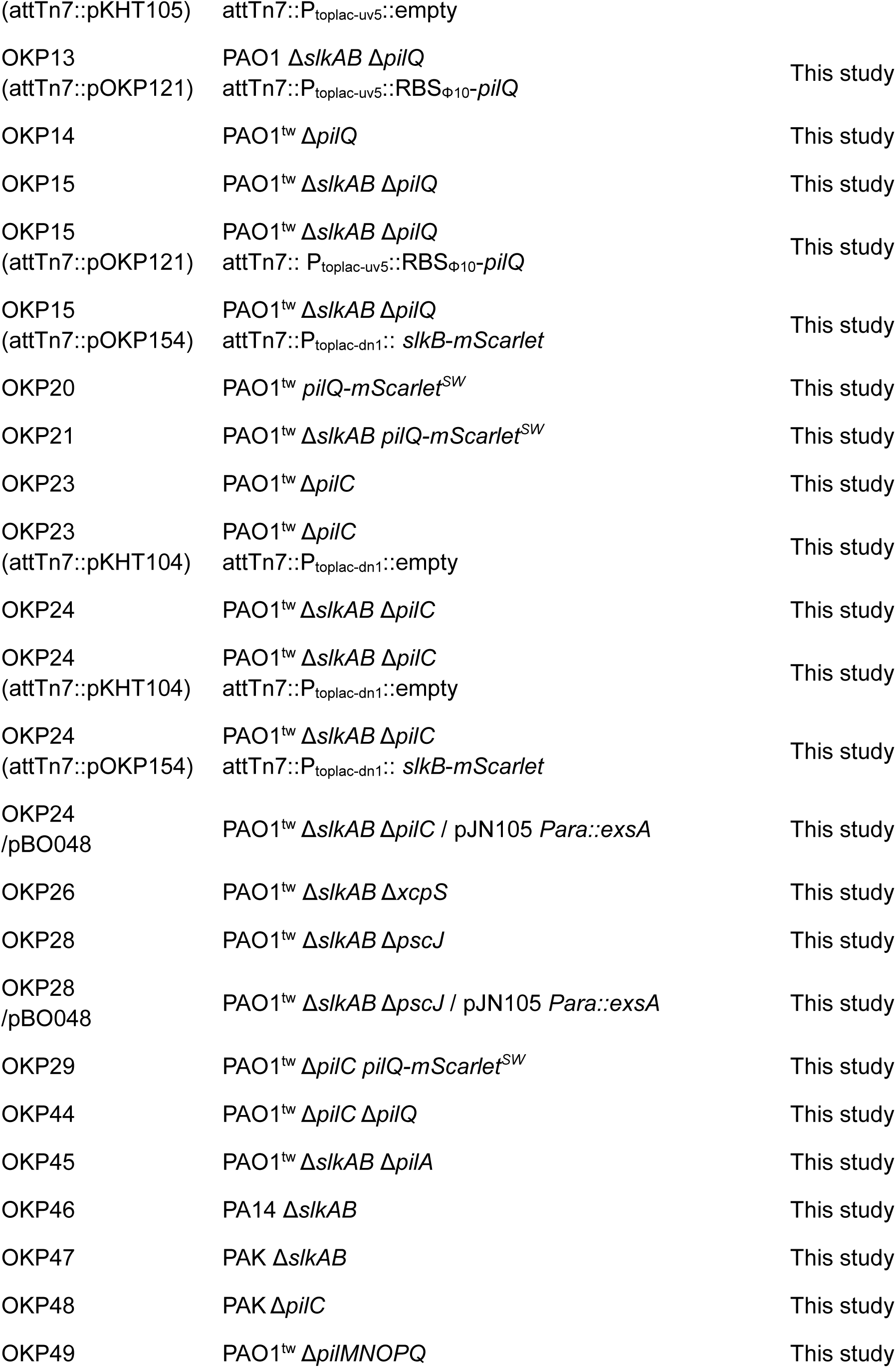
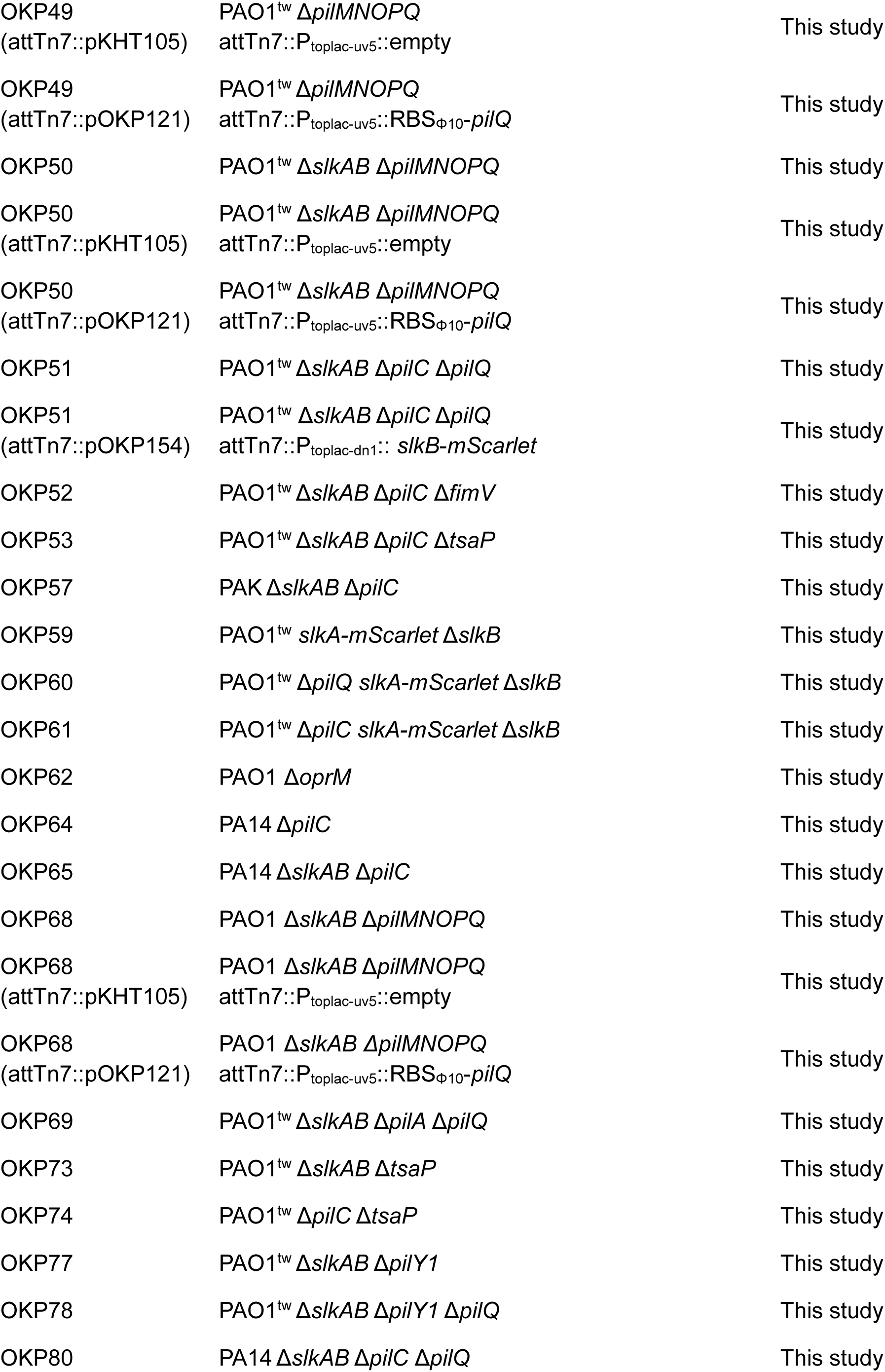
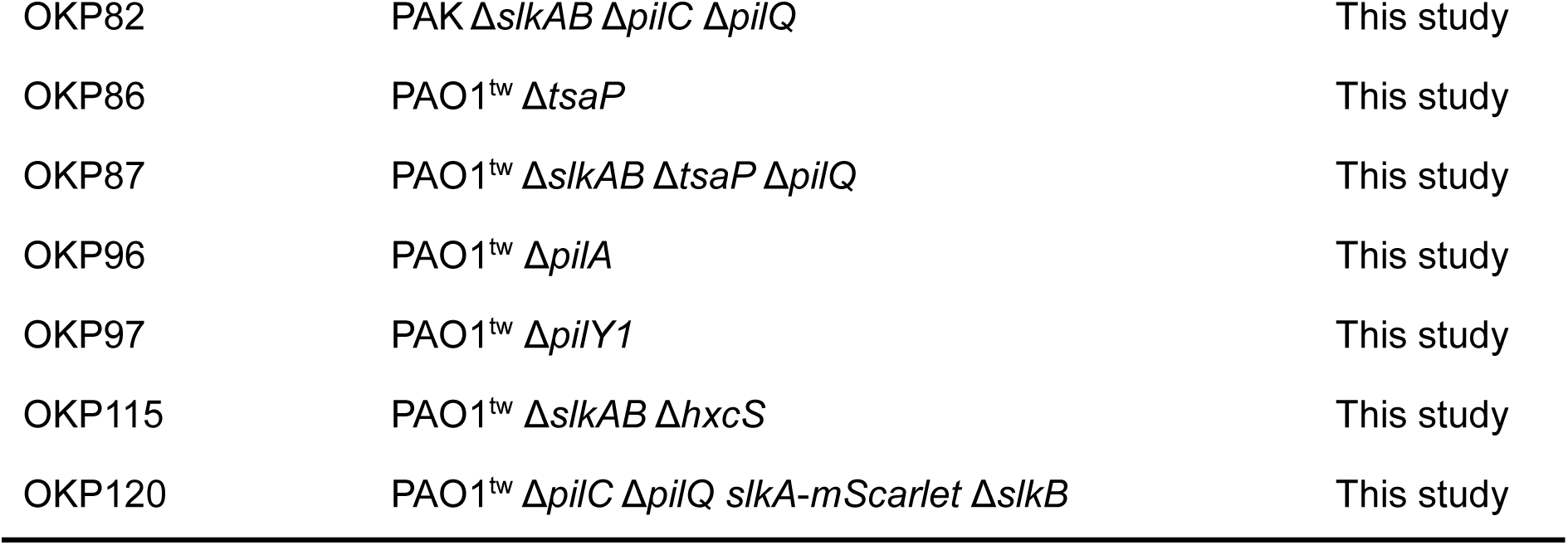
Strains used in this study.

**Table S4.**
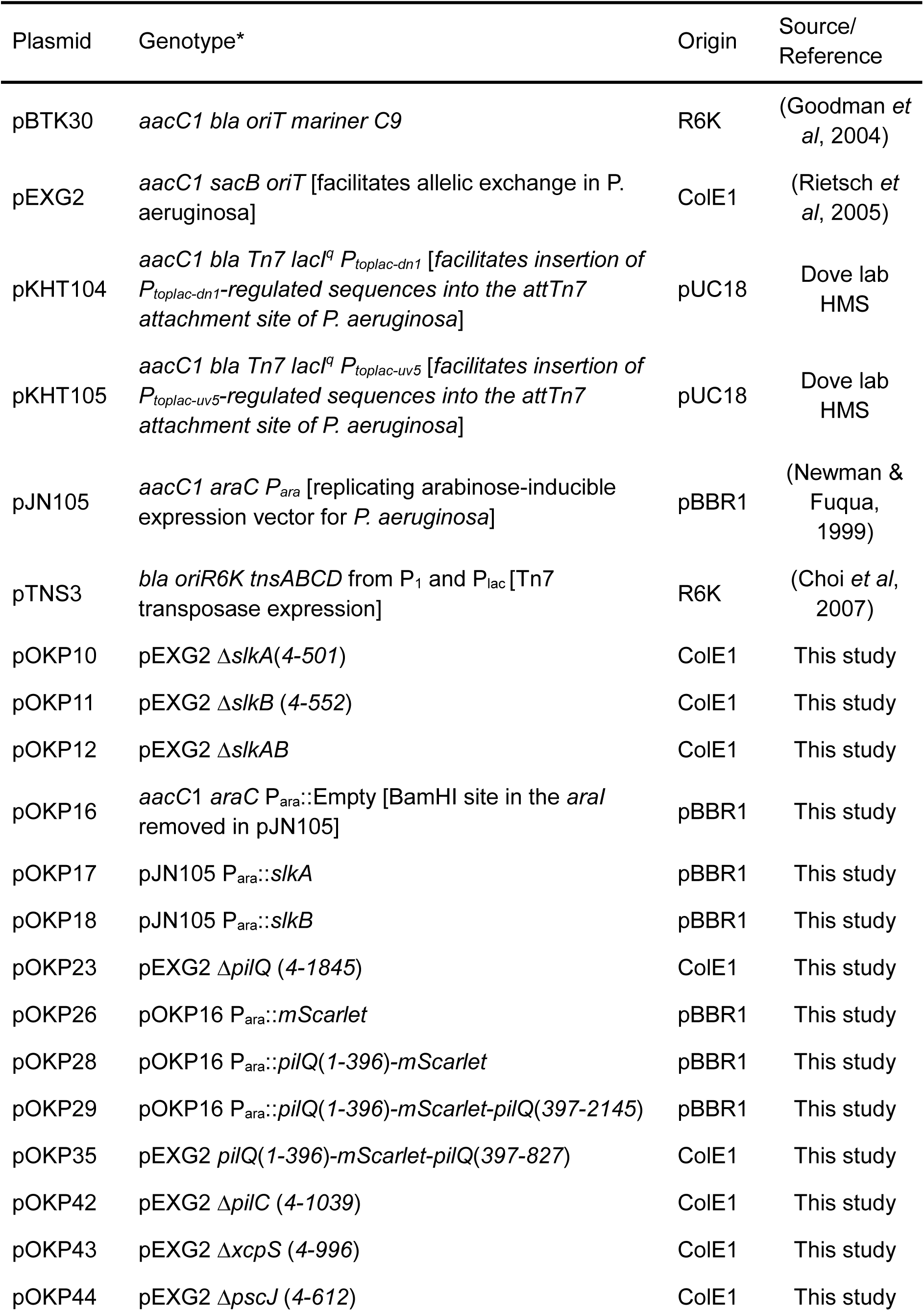
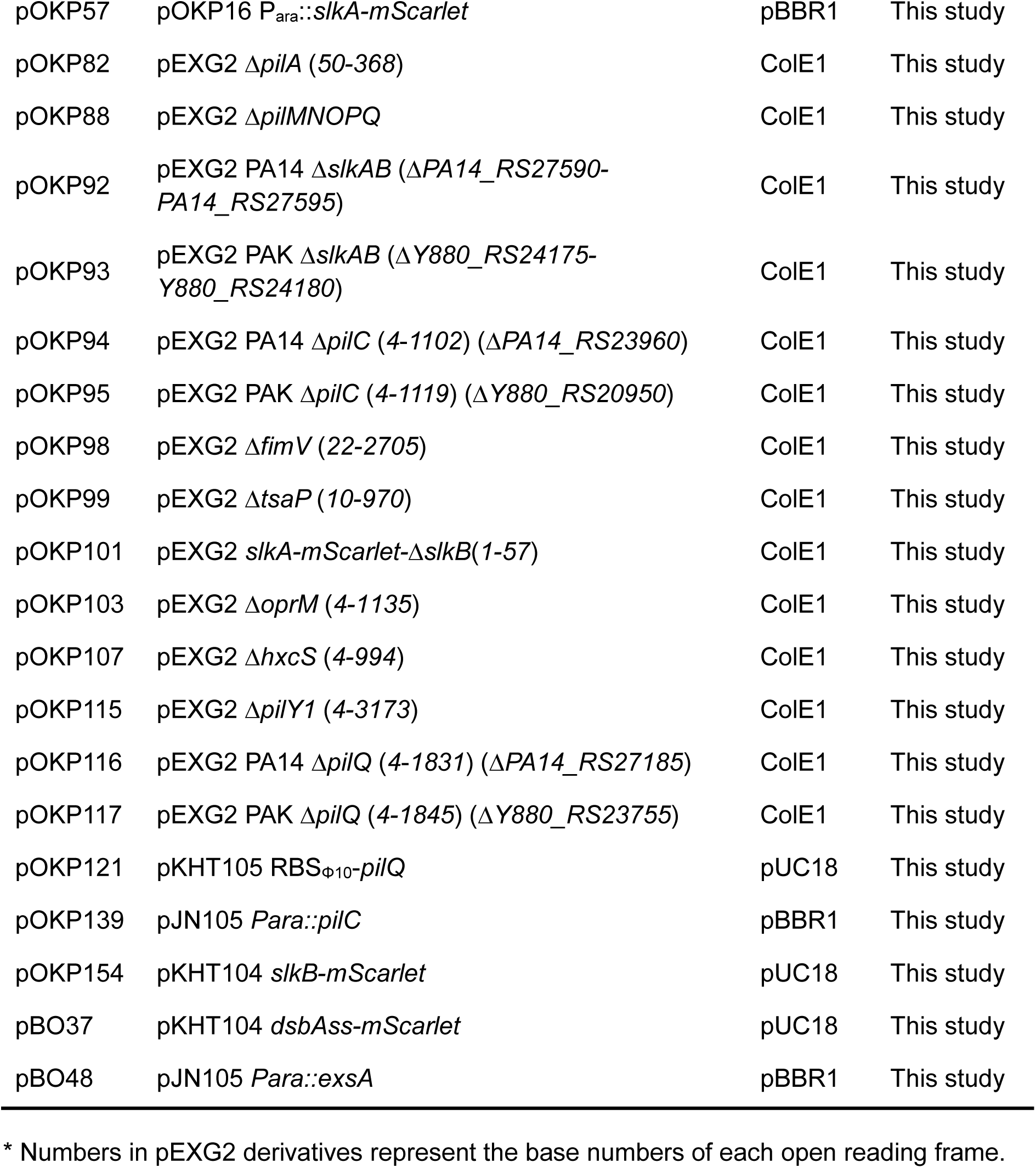
Plasmids used in this study.

